# IL-1-activated cancer-associated fibroblasts promote STAT1-driven transcriptional reprogramming of pancreatic tumour cells

**DOI:** 10.64898/2026.05.26.726819

**Authors:** Catarina Pelicano, Igor Chernukhin, Marleen Wölke, Priscilla S. W. Cheng, Lisa Young, Abigail R. Edwards, Elizabeth Mannion, Yi Cheng, Maureen Crowshaw, Steve Kupczak, Sara Pinto Teles, Muntadher Jihad, Kamal Kishore, Chandra Sekhar Reddy Chilamakuri, Valar Nila Roamio Franklin, Evangelia K. Papachristou, Clive D’Santos, Barbara Grünwald, Alasdair Russell, Jason S. Carroll, Giulia Biffi, Shalini V. Rao

**Author notes:** These authors contributed equally to this work (Igor Chernukhin, Marleen Wölke, Priscilla S. W. Cheng).

## Abstract

Pancreatic ductal adenocarcinoma (PDAC) remains one of the deadliest cancers, with limited treatment options and poor survival rates. It is characterised by strong driver mutations, epigenetic reprogramming, and a dense tumour microenvironment (TME). A defining feature of the PDAC TME is its fibrotic stroma, which is largely composed of cancer-associated fibroblasts (CAFs). Distinct CAF populations have been implicated in PDAC progression, but the mechanisms that govern their crosstalk with the cancer cells are poorly understood.

We generated genetically engineered pancreatic stellate cells (PSCs) modelling interleukin-1 (IL-1)-dependent inflammatory CAFs (iCAFs) and transforming growth factor-β (TGF-β)-dependent myofibroblastic CAFs (myCAFs) to investigate how distinct stromal populations shape the epigenetic landscape of pancreatic ductal adenocarcinoma (PDAC). We found that iCAFs, but not myCAFs, promoted gemcitabine resistance in epithelial tumour cells and identified STAT1 as a critical mediator of iCAF–tumour cell crosstalk. Mechanistically, STAT1 drove the induction of interferon (IFN)-responsive genes, while blockade of IFN-β attenuated iCAF-mediated transcriptional reprogramming. Genetic ablation of STAT1 in tumour cells abolished iCAF-induced chemoresistance and associated transcriptional changes. In an orthotopic *in vivo* model, STAT1 knockout significantly prolonged survival following gemcitabine treatment, supporting a central role for STAT1 signalling in stromal-driven therapy resistance.

We provide a comprehensive analysis on how IL-1-dependent iCAFs contribute to epigenetic reprogramming in PDAC and uncover a previously undescribed role for STAT1 in stromal-epithelial interactions. These findings reveal distinct, non-overlapping mechanisms by which CAF subtypes modulate tumour behaviour and identify STAT1 as a therapeutic vulnerability that can be exploited to sensitise PDAC to standard chemotherapy.

**Significance statement:** This study establishes that the epigenetic landscape of PDAC is differentially shaped by iCAF- and myCAF-like PSCs and defines STAT1 as a mediator of iCAF-induced chemoresistance and transcriptional reprogramming. We demonstrate that genetic ablation of STAT1 sensitises tumours to gemcitabine *in vivo*, extending survival and positioning STAT1 as an actionable target to overcome stromal-mediated therapy resistance in PDAC.

## Introduction

Pancreatic ductal adenocarcinoma (PDAC) has a median 5-year survival rate of around 10%^1,2^. Most tumours harbour a KRAS mutation that initiates a carcinogenic cascade in which cells progressively acquire dysplastic features and accumulate genetic mutations in tumour suppressor genes such as *TP53*, *CDKN2A* and *SMAD4*. Transcriptional analysis of primary PDAC tumours from several studies defined two main subtypes termed classical and squamous^3–5^. Epigenetic factors have been gradually recognised as important players in pancreatic cancer progression and as potential therapeutic targets^6–8^. Several transcription factors that orchestrate lineage differentiation during pancreas organogenesis become dysregulated in a cancer context. This is evidenced by the fact that classification between classical and squamous subtypes relies on the expression levels of a set of markers that include lineage-defining transcription factors, such as GATA, HNF and FOX proteins^4,5,9,10^. We and others have previously shown that FOXA1 promotes PDAC metastasis through enhancer reprogramming^11–13^.

PDAC is characterised by an abundant stroma mainly composed of cancer-associated fibroblasts (CAFs) and extracellular matrix (ECM) deposition. Previous work on subtyping CAF subsets defined two important subsets named inflammatory (iCAFs) and myofibroblastic CAFs (myCAFs)^14–16^. Pancreatic stellate cells (PSCs), which are a precursor population of CAFs in PDAC^17^, have been extensively used in experimental models to investigate CAF heterogeneity, and have demonstrated IL-1 signalling to promote the development of iCAFs, and TGF-β to drive differentiation toward the myCAF state^15^. Recent studies highlight the complex heterogeneity of PDAC stromal populations, and the full extent of CAF diversity in PDAC and other cancers remains unclear^18–21^. However, the functions of specific CAF-derived signals and the molecular mechanisms through which they influence epithelial cell behaviour are largely unknown. The mechanisms by which tumour microenvironment populations remodel the epithelial cell epigenome to drive tumour progression remain poorly understood. For instance, evidence suggests conditioned media from stromal populations to induce histone acetylation in pancreatic cancer cells, but the involvement of specific microenvironment populations remains unclear^22^. Here, we examine how distinct CAF subsets shape epigenetic programs in PDAC using genetically-modified CAF precursors. We show that an IL-1-driven CAF population promotes IFN-β-mediated activation of STAT1 signalling in KPC-derived cancer cells. Blocking IFN-β or deletion of *Stat1* impaired iCAF-induced transcriptional reprogramming *in vitro*, revealing a potential strategy to interfere with tumour-stroma crosstalk. Importantly, STAT1 loss significantly improved survival following gemcitabine treatment *in vivo*, implicating STAT1 in therapy resistance and highlighting this axis as a potential therapeutic target in PDAC.

## Results

### PSC-derived genetically engineered cell lines iCAF and myCAF differentially shape the epigenetic landscape of epithelial cells

The high plasticity of cancer-associated fibroblasts has made it difficult to study their function and mechanisms of action. Under standard *in vitro* culture conditions, iCAF and myCAF states are not stably maintained. In particular, fibroblasts cultured in conventional monolayer conditions predominantly acquire myofibroblastic features, whereas the inflammatory iCAF phenotype is typically induced only under specific microenvironmental cues, such as cytokine stimulation or three-dimensional co-culture systems^14^, underscoring the need to establish an experimental model that more faithfully recapitulates stable iCAF and myCAF phenotypes *in vitro* without interconversion. This limitation is further accentuated by recent work demonstrating that perturbation of TGF-β and IL-1 signalling axes can also drive repolarisation of fibroblasts into distinct CAF states *in vivo*^23^, emphasising the dynamic and signal-dependent nature of CAF identity and the challenge of faithfully modelling these phenotypes both *in vitro* and in mouse models.

To address this, pancreatic stellate cell (PSC) lines derived from the pancreata of C57BL/6J mice were genetically modified to stably reproduce and stabilise iCAF and myCAF phenotypes. Specifically, iCAF-like PSCs were generated by overexpressing IL-1α in Tgfbr2 KO PSCs^24^, whereas myCAF-like PSCs resulted from overexpressing TGF-β1 in Il1r1 KO PSCs^15^ (Fig. 1a). Protein knockout and overexpression efficiency were confirmed by Western blot and ELISA, respectively, in five lines per type (Fig. 1b and c; Extended Fig. 1a). Their gene expression profile was assessed with bulk RNA-seq (Extended Fig. 1b). Pathway enrichment analysis revealed that the transcriptional programmes of the genetically modified PSCs cultured in monolayer recapitulated previously published iCAF and myCAF signatures^14,15,19,25,24^ (Extended Fig. 1c), indicating that each cell line was. Furthermore, the modified cell lines differentially expressed key iCAF and myCAF markers (Fig. 1d), validating the distinct CAF states. Accordingly, the genetically modified iCAF- and myCAF-like PSCs stably maintained their respective phenotypes, overcoming the longstanding limitation that primary CAFs rapidly acquire myofibroblastic-dominant features under 2D culture conditions. These cell lines are therefore effectively “locked” into either an iCAF or myCAF state, providing a robust and stable model to investigate the characteristics and functions of these fibroblast subsets within the PDAC tumour microenvironment (TME). Hereafter, these cell lines are referred to as iCAFs and myCAFs.

**Figure 1.**
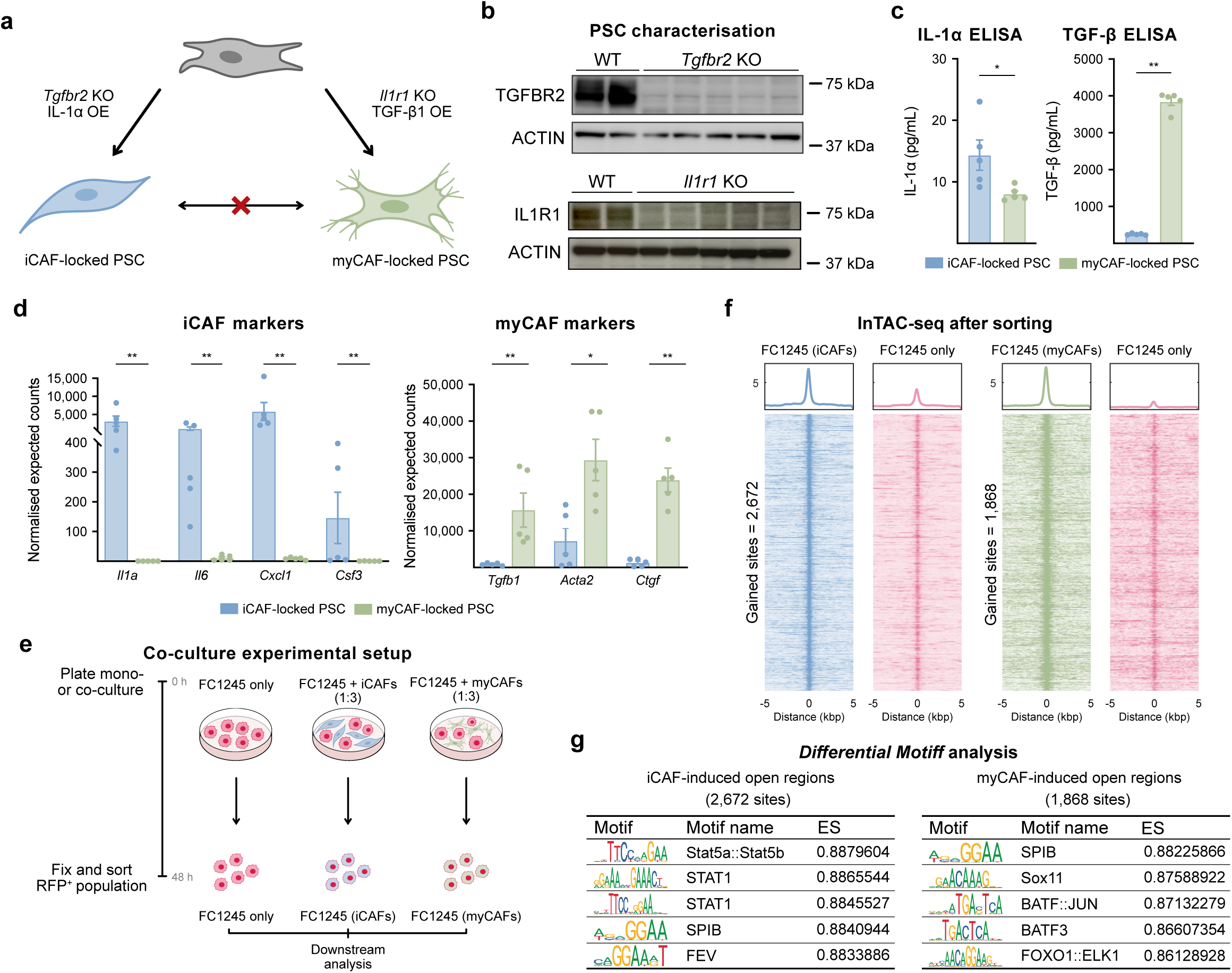
CAF-locked PSC lines shape the epigenetic state of KPC-derived epithelial cells. **a.** Schematic of genetic engineering of pancreatic stellate cells (PSCs) into iCAF-locked and myCAF-locked states. KO, knockout. OE, overexpression. **b.** Western blot analyses of TGFBR2 and IL1R1 in iCAF- (left) and myCAF- (right) locked PSCs, respectively. Three *Tgfbr2* gRNAs and two *Il1r1* gRNAs were used for CRISPR knockout, respectively. Parental PSCs were used as positive controls. ACTIN was used as loading control. **c.** Normalised expected counts of iCAF (left panel; *Il1a*, *Il6*, *Cxcl1* and *Csf3*) and myCAF (right panel; *Tgfb1*, *Acta2* and *Ctgf*) markers in iCAF-locked and myCAF-locked PSCs. Results show mean ± SEM. Statistical significance was calculated by Mann-Whitney U test. \**p* ≤ 0.05*; ** p* ≤ 0.01. **d.** IL-1α and TGF-β ELISA of culture media from iCAF-locked and myCAF-locked PSCs. Statistical significance was calculated by Mann-Whitney U test. \**p* ≤ 0.05; *\*\* p* ≤ 0.01. **e.** Schematic of the co-culture experimental setup. **f.** *DiffBind*^27^ heatmaps of ATAC-seq signal at open regions gained in FC1245-RFP^+^ after co-culture with iCAFs (left panel) and FC1245-RFP^+^ after co-culture with myCAFs (right panel) compared FC1245-RFP^+^ alone, n=4. **g.** Top differential motifs of newly open regions induced by co-culture with iCAFs (top panel) and myCAFs (bottom panel). ES, enrichment score.

To determine whether the epigenetic landscape of tumour cells was influenced by the presence of these CAF populations, KPC (*Kras*^G12D/+^; *Trp53*^R172H/+^; *Pdx1*-Cre) mice-derived FC1245-RFP^+^ cells were cultured alone or together with iCAFs or myCAFs, and then sorted for downstream analysis (Fig. 1e). To investigate whether iCAF- and myCAF-“locked” lines influenced chromatin dynamics of PDAC cells, we adapted a modified assay for transposase-accessible chromatin using sequencing (ATAC-seq) protocol described by Baskar *et al.*^26^ to profile chromatin accessibility from fixed, sorted cells. *DiffBind*^27^ analysis revealed differences in chromatin accessibility between cells that had been cultured alone and cells that had been co-cultured with either CAF population. Specifically, iCAFs induced the opening of 2,672 genomic regions, whereas myCAFs increased the accessibility of 1,868 sites (Fig. 1f). DiffMotif analysis further indicated distinct enriched motifs: while regions open after iCAF co-culture were mostly enriched in STAT motifs, regions opened by myCAFs were enriched in SOX and JUN motifs, among others (Fig. 1g), suggesting that these two populations could play different roles in shaping the epigenetic landscape of tumour cells.

### iCAFs but not myCAFs promote chemoresistance in cancer cells

To assess whether the chromatin accessibility changes resulted in differential gene expression, we performed RNA-seq on co-cultures (Fig. 1e), which revealed distinct transcriptomic profiles for FC1245-RFP^+^ cells after co-culture with either CAF populations (Fig. 2a). GSE analysis showed that iCAFs upregulated pathways such as “IFN-α response” and “IFN-γ response” in cancer cells, whereas myCAFs drastically upregulated “epithelial-to-mesenchymal transition”, as well as “apical junction”, “TNF-α signalling via NF-κB” and “hypoxia” transcriptional programmes (Fig. 2b). These pathways differed from those stimulated by non-genetically locked PSCs^22^, suggesting that IL-1 and TGF-β activated CAFs could play distinct, specific roles in pancreatic tumours.

**Figure 2.**
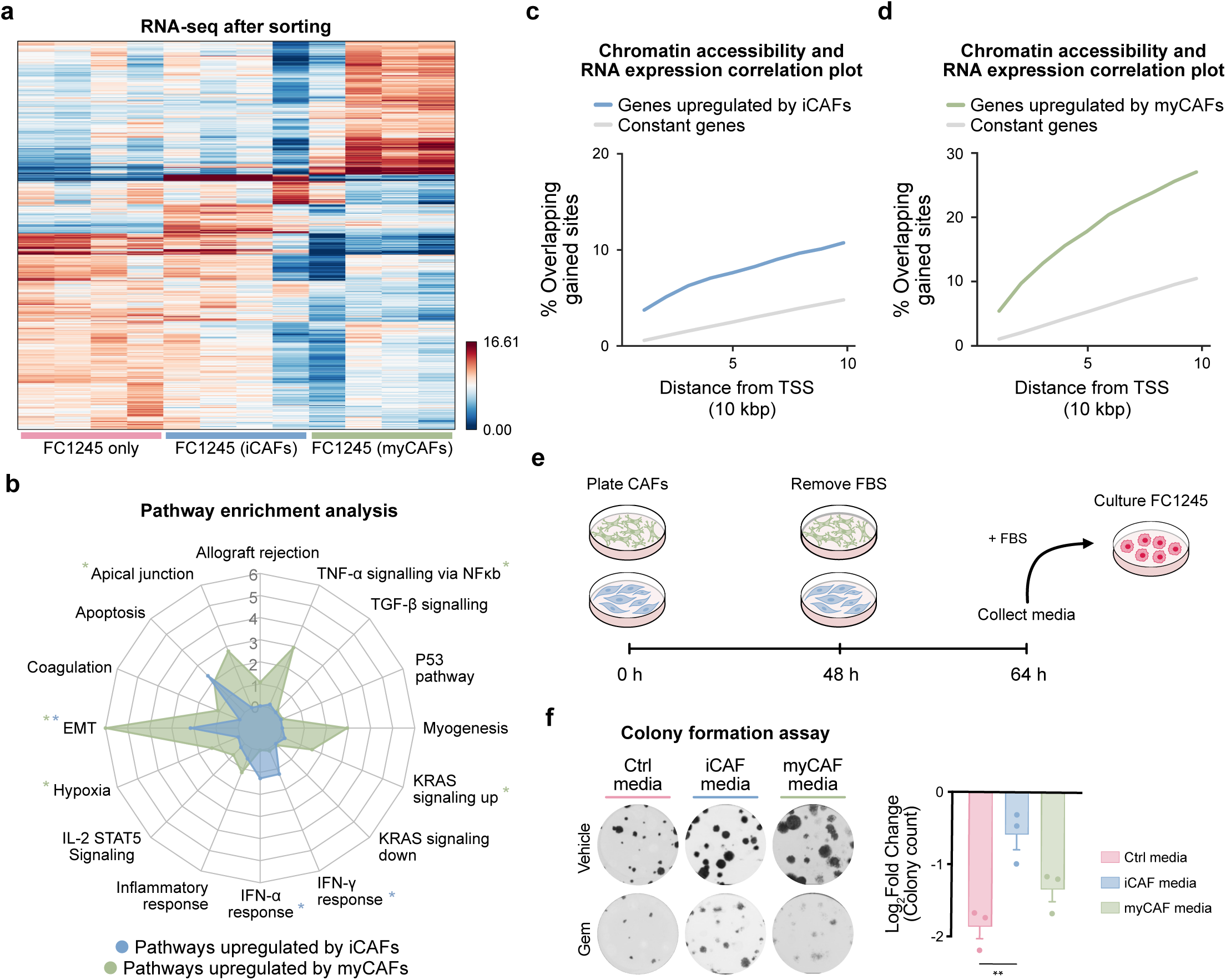
CAF-locked PSC lines influence transcriptional programs of epithelial cells. **a.** Differential gene expression analysis of FC1245-RFP^+^ cells cultured alone or sorted from co-culture with iCAFs or myCAFs, n=4. **b.** Spider plot showing GSEA pathway enrichment after co-culture with iCAFs or myCAFs compared to control. Plotted values correspond to enrichment scores for the comparisons FC1245 (iCAFs) vs FC1245 only (blue) and FC1245 (myCAFs) vs FC1245 only (green). Pathways significantly enriched after iCAF or myCAF co-culture are marked with a blue or green asterisk, respectively. Statistical significance was calculated using the GSEA-based Kolmogorov-Smirnov statistical test, \**p* < 0.05. **c.** Enrichment of newly accessible regions in the vicinity of upregulated genes compared to constitutively expressed genes in FC1245-RFP^+^ cells after co-culture with iCAFs. **d.** Enrichment of newly accessible regions in the vicinity of upregulated genes compared to constitutively expressed genes in FC1245-RFP^+^ cells after co-culture with myCAFs. **e.** Schematic representation of the experimental setup used for conditioned media experiments. **f.** Colony formation assay of FC1245-RFP^+^ cells cultured in control, iCAF- or myCAF-conditioned media and treated with 5 nM of gemcitabine for 7 days. Results are relative to vehicle (DMSO), n=3. Statistical significance was calculated using a one-way ANOVA, \*\**p*=0.0046.

To investigate whether the newly open chromatin regions were responsible for the differential gene expression observed, we assessed the correlation between the two datasets. Indeed, regions that gained accessibility upon co-culture with iCAFs (2,672 sites) were enriched in the vicinity of upregulated genes (Fig. 2c). Similarly, myCAF-induced newly open regions (1,868 sites) were in closer proximity to genes upregulated after co-culture (Fig. 2d). This correlation indicated that regions with increased chromatin accessibility were in part responsible for the transcriptional reprogramming observed in each context, supporting the notion that the CAFs mediate meaningful epigenetic changes in pancreatic cancer cells.

Since IFN-response genes have been associated with increased resistance to chemotherapy agents^28,29^ and this pathway was upregulated after co-culture with iCAFs, we sought to test whether cytokines released by the CAF lines and upstream of IFN signalling could exert a similar effect in response to gemcitabine. To investigate this, FC1245-RFP^+^ cells were cultured using conditioned media derived from iCAFs or myCAFs as detailed in Fig. 2e. Cancer cells cultured in iCAF-but not myCAF-conditioned media showed a significant increase in chemoresistance to gemcitabine (Fig. 2f).

### iCAFs and myCAFs accelerate tumour progression *in vivo* without affecting FOXA1

We sought to assess whether the CAF subsets could modulate cancer progression *in vivo*. To test this, FC1245-RFP^+^ cells were pre-cultured alone or together with iCAFs-GFP^+^ or myCAFs-GFP^+^ prior to flow sorting (Extended Fig. 2a). Subsequently, these cells were orthotopically injected into murine pancreases (Fig. 3a). Mice that were co-injected with tumour cells and either or both subsets of fibroblasts showed reduced survival compared to mice injected with tumour cells only (Fig. 3b), suggesting that both these CAF subtypes are pro-tumorigenic. However, postmortem immunohistochemistry analysis of the tumours revealed low to zero GFP staining in all cohorts (Extended Fig. 2b). While these data may be consistent with a limited contribution of the *in vivo* interactions to the observed survival differences, they do not exclude other hypotheses, for instance, the possibility of the locked lines influencing cancer cells at earlier stages of tumour implantation before being lost. Furthermore, the co-culture step prior to injection could impart a priming effect or enhance engraftment. This suggests an important role for CAF-mediated epigenetic reprogramming in this context that remains to be further explored.

**Figure 3.**
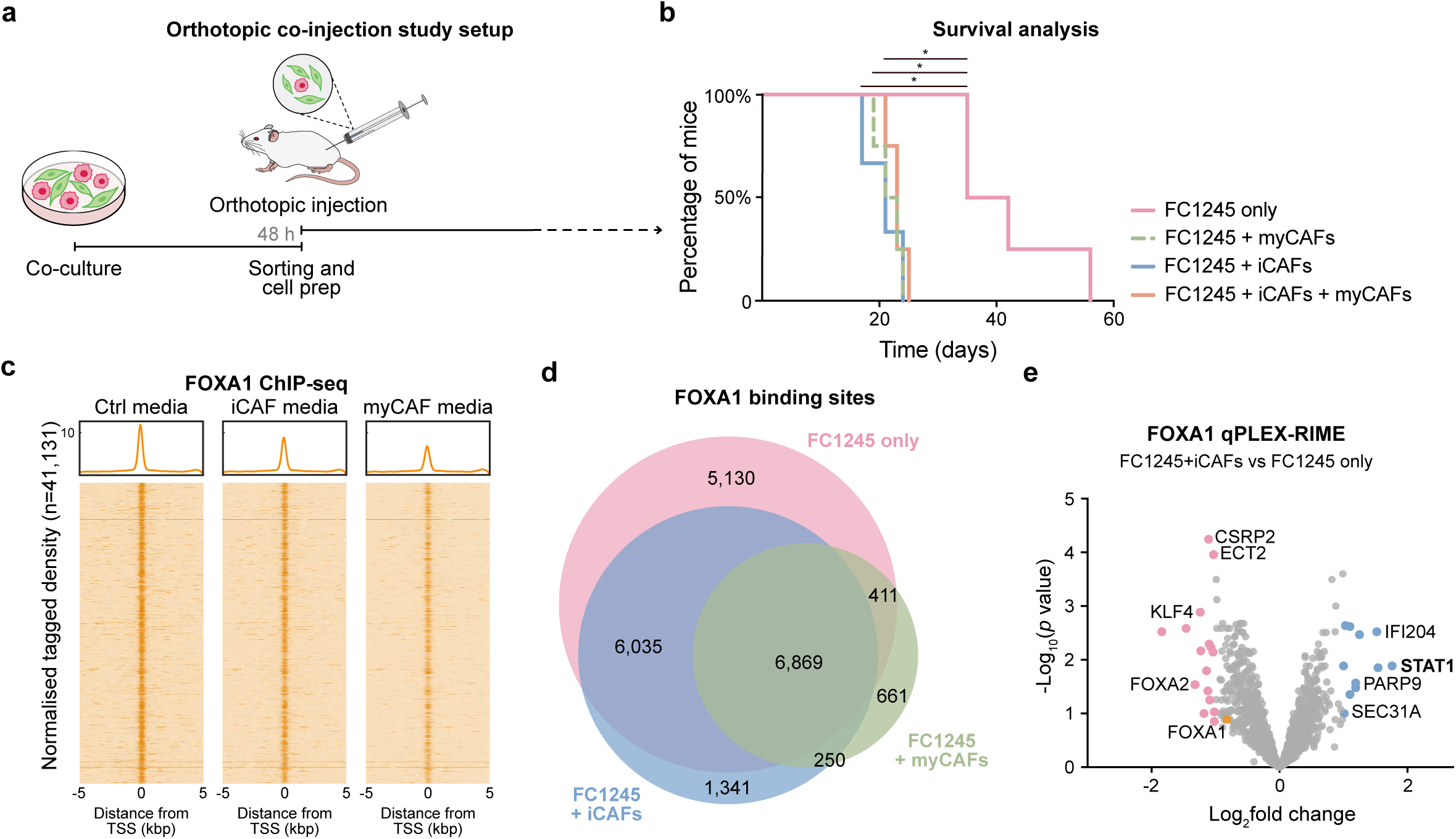
iCAFs and myCAFs promote tumour progression *in vivo*. **a.** Schematic representation of the co-injection *in vivo* study timeline. In summary, FC1245-RFP^+^ cells were cultured alone or together with iCAFs-GFP^+^ or myCAFs-GFP^+^ for 48 hours, after which cells were FACS-sorted and mixed at a 1:4 ratio. Cells were resuspended in 50 % Matrigel in PBS and orthotopically injected onto the pancreas of the mice. **b.** Kaplan-Meier survival plots of mice orthotopically injected with FC1245-RFP^+^ cells alone or together with iCAFs and/or myCAFs (1:4 ratio), n=4. Statistical significance was calculated using a log-rank (Mantel-Cox) test with a Bonferroni multiple comparison correction, \**p* < 0.05. **c.** Normalised tagged intensity heatmaps of FOXA1 ChIP-seq signal in FC1245-RFP^+^ cultured alone or together with iCAFs or myCAFs, n=3. **d.** Venn diagram indicating the overlap of FOXA1 binding sites across conditions. **e.** Volcano plot of FOXA1 qPLEX-RIME data representing FOXA1 interactors differentially regulated after co-culture with iCAFs, n=6. Proteins with log_2_fold change > 1 and adj. *p* value < 0.05 are highlighted in blue (significantly upregulated by iCAFs) and proteins with log_2_fold change < -1 and adj. *p* value < 0.05 are highlighted in pink (significantly upregulated in monoculture). Statistical significance was calculated using a limma-based differential analysis.

We and others have previously highlighted the role of the pioneer factor FOXA1 in pancreatic cancer metastasis^11–13^. However, the influence of the tumour microenvironment in regulating FOXA1’s function remains unexplored. As both CAF populations significantly accelerated disease progression *in vivo* (Fig. 3b) and upregulated EMT pathway in tumour cells *in vitro* (Fig. 2b), we hypothesised that FOXA1 could bind regulatory elements that promote tumour growth downstream of CAF-epithelial cell crosstalk.

To test this, we performed FOXA1 chromatin immunoprecipitation followed by sequencing (ChIP-seq) on FC1245-RFP^+^ cultured alone or together with iCAFs or myCAFs, which demonstrated a reduction in total FOXA1 binding sites and intensity after co-culture with either iCAFs or myCAFs compared to control (Fig. 3c and d). Furthermore, the genomic distribution of FOXA1 remained unchanged across conditions (Extended Fig. 2c), indicative of reduced FOXA1 occupancy without evidence of large-scale reprogramming. These data indicate that both co-culture settings broadly reduced FOXA1 binding intensity without altering its underlying genomic binding, suggesting that tumour-promoting CAF signals might be acting independently of FOXA1 regulatory elements in this context. To further dissect how each CAF population influences the FOXA1 interactome, we then performed qPLEX-RIME^30,31^, a quantitative approach to profile protein-protein interactions on chromatin. Differential interactome analysis revealed the recruitment of STAT1 to the FOXA1 complex upon co-culture with iCAFs (Fig. 3e), but not with myCAFs (Extended Fig. 2d). Notably, STAT1 and FOXA1 have previously been reported to interact at the chromatin level in prostate and breast cancer models^32^. Given that the iCAFs also selectively induced upregulation of IFN-α and IFN-γ response pathways in cancer cells (Fig. 2b) and increased accessibility of chromatin regions enriched in STAT motifs (Fig. 1g), we focused subsequent analyses on STAT1-dependent mechanisms underlying iCAF-cancer cell crosstalk.

### STAT1 is recruited to chromatin of cancer cells in response to iCAF-derived cues

To further confirm STAT1 pathway activation following iCAF co-culture, we assessed *Stat1* expression levels in our RNA-seq dataset (Fig. 2b) and performed a Western blot to evaluate STAT1 protein levels before and after co-culture. Both STAT1 mRNA and protein levels were elevated after co-culture with iCAFs, but not with myCAFs, compared to the control (Fig. 4a and 4b), further suggesting a role for this transcription factor downstream of iCAF-derived signals specifically. Indeed, STAT1 was upregulated and phosphorylated at both Tyrosine 701 and Serine 272 residues after co-culture with iCAFs, but not myCAFs (Extended Fig. 3a). Consistent with previous findings^15^, STAT1 protein levels were also higher in the iCAF population compared to the myCAFs (Fig. 4b).

**Figure 4.**
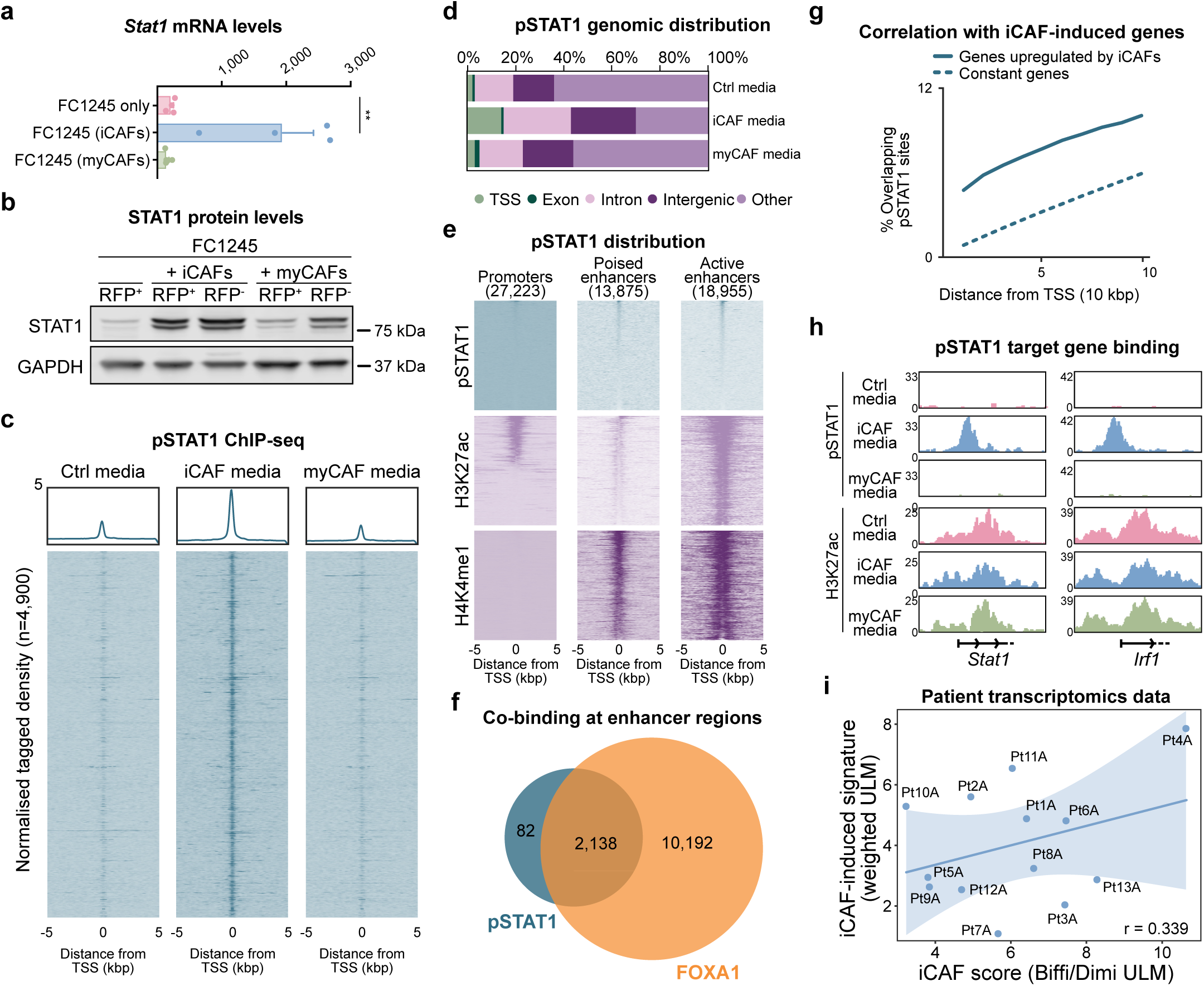
iCAFs but not myCAFs promote the recruitment of STAT1 to chromatin. **a.** *Stat1* mRNA levels in FC1245-RFP^+^ cells cultured alone or sorted from co-culture with iCAFs or myCAFs. Statistical significance was calculated using a one-way ANOVA, \*\**p*=0.003. **b.** Western blotting of STAT1 in FC1245 (RFP^+^) or CAF populations (RFP^-^) after co-culture for 48 hours (left panel). **c.** Normalised tagged intensity heatmaps of pSTAT1 ChIP-seq signal in FC1245-RFP^+^ cells cultured in control, iCAF- or myCAF-conditioned media, n=4. **d.** Genomic feature distribution of pSTAT1 in FC1245-RFP^+^ cells cultured in iCAF-conditioned media compared to control media. **e.** pSTAT1 binding distribution across promoter (H3K27ac high, H3K4me1 low), poised enhancer (H3K27ac low, H3K4me1 high) and active enhancer (H3K27ac high, H3K4me1 high) regions. **f.** Venn diagram indicating the overlap between pSTAT1^Tyr701^ and FOXA1 binding at enhancer regions (H3K4me1^hi^). FC1245-RFP^+^ cells cultured in iCAF-conditioned media or together with iCAFs, respectively. **g.** Enrichment of pSTAT1 binding sites (upon iCAF media) in the vicinity of genes upregulated after iCAF co-culture compared with constitutively expressed genes in FC1245-RFP^+^ cells. **h.** Representative IGV tracks of pSTAT1 and H3K27ac binding at the *Stat1* and *Irf1* promoters across conditions. **i.** Scatter plot showing the correlation between iCAF-induced gene signature scores (weighted ULM) and iCAF signature scores (ULM, indicating iCAF presence) across spatial transcriptomic datasets of PDAC patients from Pei *et al*^34^. Each dot represents the median of all spots from each individual patient. The iCAF scores were quantified using previously established iCAF signature from Biffi *et al.*^15^, and the iCAF-induced signature scores were calculated using genes upregulated in FC1245-RFP^+^ cells after co-culture with iCAFs.

To investigate pSTAT1 chromatin binding, we used the conditioned media experimental setup as described above (Fig. 2e). ChIP-seq analysis confirmed that pSTAT1 binding at functional genomic regions occurred almost exclusively upon stimulation with iCAF-conditioned media, and not when cells were cultured in control or myCAF-conditioned media (Fig. 4c and d and Extended Fig. 3b). pSTAT1 binding was also accompanied by H3K27ac, indicating association with transcriptionally active chromatin (Extended Fig. 3c). Further integration of H3K27ac and enhancer hallmark H3K4me1 profiles following iCAF co-culture revealed that pSTAT1 primarily localised to active enhancer regions (Fig. 4e), suggesting that STAT1 recruitment occurs at pre-existing regulatory elements that become further activated in this context. We hypothesised that these regions could be pre-bound by FOXA1, since more than half of the total pSTAT1 binding sites overlapped with FOXA1 baseline sites found across conditions, independently of CAF cues (Extended Fig. 3d). Notably, most of the pSTAT1-bound enhancer regions (marked by H3K4me1) were also co-bound by FOXA1 (Fig. 4f). This suggests that FOXA1 pre-marks a subset of potentially poised regulatory elements prior to pSTAT1 recruitment in response to upstream signalling cues. These findings support a model in which iCAF-derived signals contribute to enhancer activation at FOXA1-bound, interferon-responsive loci. Furthermore, pSTAT1 chromatin occupancy following stimulation with iCAF-conditioned media positively correlated with gene expression changes observed after iCAF co-culture (Fig. 4g), indicating that pSTAT1 binding sites play an important role in regulating transcriptional reprogramming downstream of iCAF-derived signals. Target genes included, for instance, *Stat1* and *Irf1* (Fig. 4h), consistent with previously published datasets^33^. Overall, these data position STAT1 as a crucial downstream effector of iCAF-mediated signalling.

Having established that iCAF-regulated transcriptional reprogramming is partially driven by STAT1, we derived an iCAF-induced transcriptional signature based on our co-culture data (Fig. 2a) and sought to validate its association with iCAFs in primary tumours from PDAC patients. Using spatial transcriptomic datasets from Pei *et al.*^34^, we found that iCAF enrichment scores positively correlated with the iCAF-induced signature derived from our model (Fig. 4i and Extended Fig. 3e). These data suggests that the transcriptional changes observed *in vitro* recapitulate features of the tumour microenvironment of PDAC patients.

### IFN-***β*** blockade or STAT1 KO impair iCAF-mediated transcriptional reprogramming

To further assess the transcriptional pathways regulated by pSTAT1 in the presence of iCAFs, GSEA was conducted on genes located downstream of pSTAT1-bound promoters. As expected, this analysis revealed enrichment of IFN-α and IFN-γ response pathways (Extended Fig. 4a), which indicated that pSTAT1 is in part responsible for the iCAF-induced transcriptional reprogramming (Fig. 2b). Given these results, we hypothesised that an interferon signalling molecule would be upstream of STAT1 phosphorylation and constitute a key mediator of the iCAF-epithelial cell crosstalk. Blocking IFN-β, but not IFN-α or IFN-γ, significantly reduced STAT1 phosphorylation levels (Fig. 5a and Extended Fig. 4b) and decreased the gene expression changes observed upon stimulation with iCAF media (Fig. 5b), suggesting a way to interfere with this crosstalk. Indeed, the genetically-locked iCAFs exhibited higher IFN-β expression levels compared to the myCAFs (Extended Fig. 4c). Blocking conventional iCAF markers such as IL-6 or IL-1α did not result in a reduction of STAT1 phosphorylation levels (Extended Fig. 4d), suggesting that these cytokines are not critical for pathway activation in this context. Our results therefore uncover a previously undescribed role for IFN-β in the iCAF-epithelial crosstalk in PDAC.

**Figure 5.**
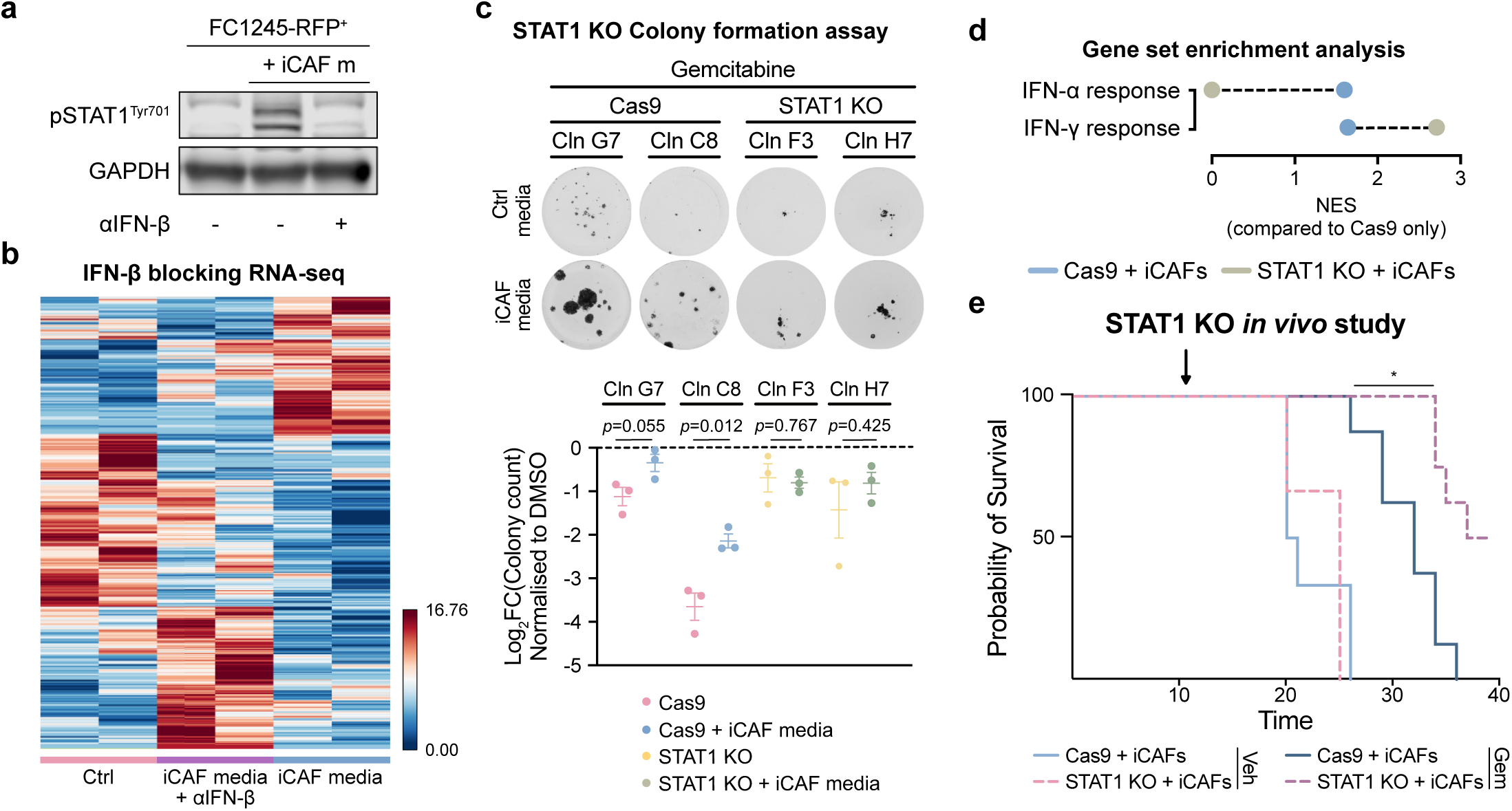
STAT1 perturbation impairs iCAF-induced transcriptional reprogramming. **a.** Western blotting of pSTAT1^Tyr701^ in FC1245-RFP^+^ cells cultured in control, iCAF-conditioned media or iCAF-conditioned media treated with 2,000 neutralising units (NU)/mL of anti-IFN-β for 48 hours. **b.** Heatmap of differential gene expression of FC1245-RFP^+^ cells cultured in control or iCAF-conditioned media supplemented or not with anti-IFN-β. **c.** Colony formation assay of STAT1 KO or Cas9 only clones cultured in control or iCAF-conditioned media and treated with 5 nM of gemcitabine. Results are relative to vehicle (DMSO), n=3. Statistical significance was calculated using a one-way ANOVA, \**p* = 0,0158. **d.** GSE analysis of gene expression patterns in Cas9 or STAT1 KO clones after co-culture with iCAFs (compared to Cas9 alone). **e.** Kaplan-Meier survival plots of mice orthotopically injected with STAT1 KO or Cas9 control clones together with iCAFs (1:10 ratio) and treated with gemcitabine or vehicle twice a week, n=6 for vehicle mice and n=8 for gemcitabine-treated mice. Arrow indicates treatment start date. Statistical significance was calculated using a log-rank (Mantel-Cox) test with a Bonferroni multiple comparison correction, \**p* < 0.05.

To investigate whether the iCAF-mediated epigenetic modulation of epithelial cells was predominantly dependent on STAT1, we generated FC1245-RFP^+^ with CRISPR-Cas9-mediated knockout of *Stat1* (STAT1 KO, Extended Fig. 5a). We first assessed colony formation capacity of Cas9 and STAT1 KO clones following gemcitabine treatment. Consistent with our previous observations with the parental line (Fig. 2f), Cas9 controls exhibited increased resistance to gemcitabine when cultured in iCAF-conditioned media (compared to control media). In contrast, STAT1 knockout clones did not exhibit this advantage (Fig. 5c), indicating that STAT1 is required for the iCAF-induced chemoprotective phenotype.

We next assessed whether STAT1 loss would abrogate the previously observed iCAF-induced changes in chromatin accessibility (Fig. 1f). To this end, two independent STAT1 knockout clones and two Cas9 control clones were pooled, and the two pools were cultured either alone or in co-culture with iCAFs, and subsequently sorted prior to ATAC-seq^26^ analysis. Notably, we observed that these regions become more accessible independent of STAT1 status following iCAF co-culture (Extended Fig. 5b), indicating that the factors responsible for chromatin opening in this context are independent or possibly upstream of STAT1.

To determine if STAT1 loss impacted the transcriptional response to iCAF-derived stimuli, we next performed RNA-seq on the pooled STAT1 knockout and Cas9 control clones before and after co-culture with iCAFs. STAT1 KO and Cas9 control samples clustered closely together, indicating that core transcriptional programmes remain largely preserved following STAT1 loss, whereas Cas9 cells exposed to iCAFs formed a distinct cluster. These findings suggest that tumour microenvironmental signals derived from iCAFs are the primary drivers of transcriptional reprogramming in this context, highlighting a critical role for STAT1 in mediating the downstream response (Extended Fig. 5c). GSE analysis confirmed enrichment of IFN-α and IFN-γ response pathways in Cas9 controls following co-culture with iCAFs (Fig. 5d), consistent with our observations in the parental line (Fig. 2b). In contrast, STAT1 knockout cells co-cultured with iCAFs showed a complete loss of IFN-α response enrichment, although IFN-γ response signatures were further elevated relative to Cas9 controls. Notably, gene signatures upregulated in the STAT1 KO clones following iCAF co-culture demonstrated reduced association with iCAF-induced open regions (Fig. 1f) compared to Cas9 controls (Extended Fig. 5d). These date indicates that gene expression downstream of iCAF signalling is reduced in the absence of STAT1. Together, our findings suggest that STAT1 is required for the iCAF-mediated transcriptional reprogramming and chemoresistance.

### STAT1 KO extends survival upon gemcitabine treatment in an orthotopic *in vivo* ***model***

We then sought to investigate whether STAT1 loss would sensitise tumours to gemcitabine treatment in an orthotopic model. Cas9 control or STAT1 knockout FC1245-RFP^+^ cells were co-injected together with iCAFs into murine pancreases, and mice were subsequently treated with either vehicle or gemcitabine. While STAT1 deletion alone did not significantly affect survival in vehicle-treated cohorts, STAT1 knockout tumours exhibited markedly improved survival compared to Cas9 controls in the treated cohorts (Fig. 5e), demonstrating that STAT1 is required for chemoresistance *in vivo*. These data indicate that STAT1 deletion in the tumour cells specifically sensitizes tumours to chemotherapy in the presence of iCAFs, rather than affecting baseline tumour growth. Together with our *in vitro* observations, these results position STAT1 as a critical therapeutic target to overcome stromal-mediated chemoresistance in pancreatic cancer.

To dissect the specific role of STAT1 in a patient context, we derived a transcriptomic signature of the genes upregulated in Cas9 controls compared to STAT1 KO in the presence of iCAFs (Cas9 + iCAFs vs STAT1 KO + iCAFs score). This signature positively correlated with the previously derived iCAF-induced signature in transcriptomic datasets of PDAC patients^34^, suggesting that the transcriptomic changes mediated by iCAFs are highly dependent on STAT1 (Fig. 6a). Furthermore, analysis of patient data from the Moffit *et al*. dataset^4^ revealed upregulation of *Stat1* mRNA in primary tumour samples compared to normal tissue (Fig. 6b), suggesting a potential role for this transcription factor in the transition to malignancy. In contrast, *Stat1* expression was reduced in metastatic samples relative to primary tumours, indicative of a stage-specific function in a primary tumour context. Analysis of untreated patients of the COMPASS trial dataset further revealed that, in late-stage disease, higher *STAT1* expression in the primary tumour is associated with poorer overall survival (Fig. 6c). Notably, this association was not observed in early-stage disease. This stage-specific effect further suggests a context-dependent role for STAT1 in PDAC progression. Given than desmoplasia and fibroblast activation become more pronounced during progression to invasive PDAC^26,27^, one possible explanation is that signalling cues from iCAFs become more prominent or functionally consequential in advanced disease, thereby contributing to enhanced STAT1 activation at later stages.

**Figure 6.**
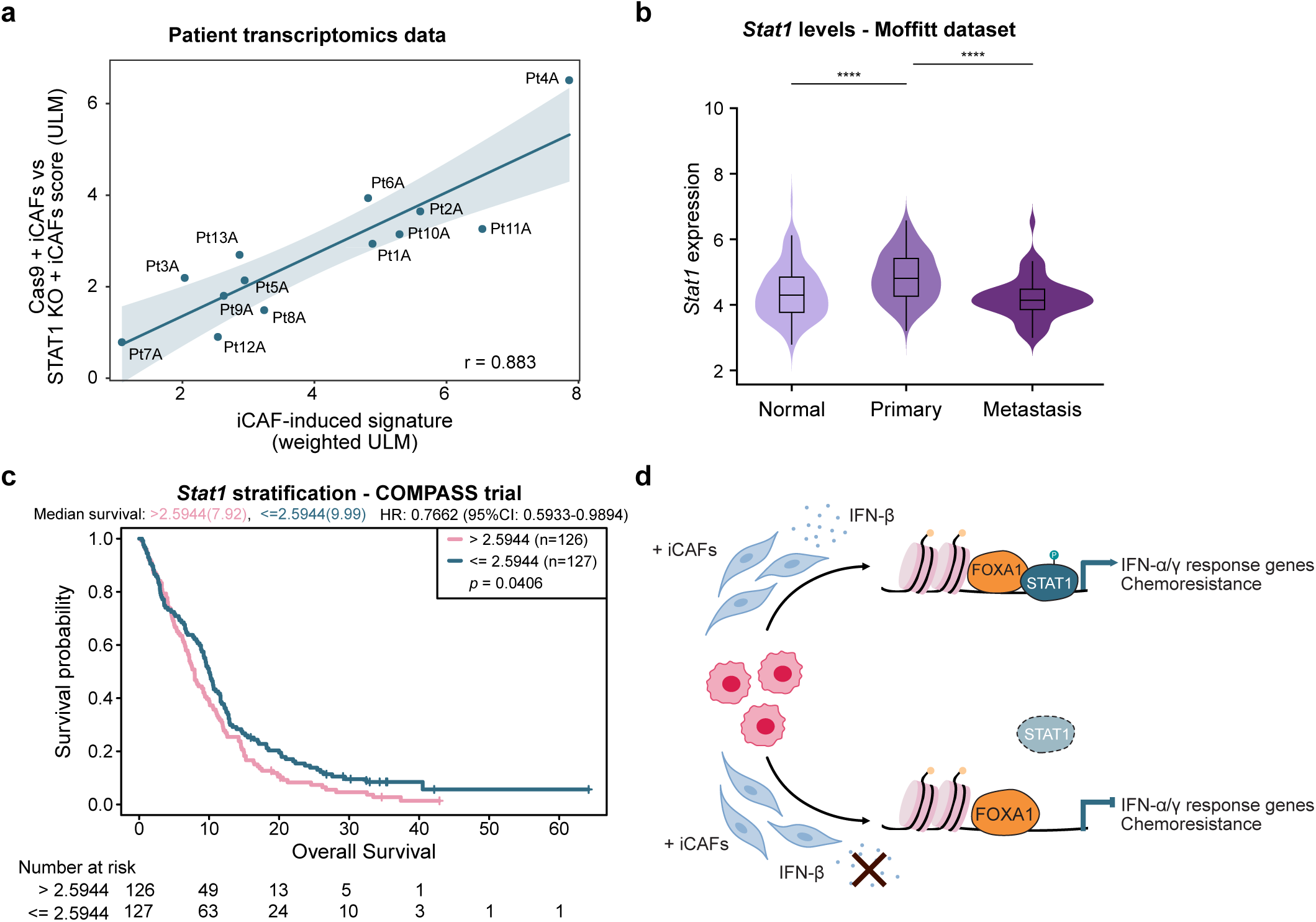
STAT1 is associated with poor prognosis in PDAC patients. **a.** Scatter plot showing the correlation between Cas9 + iCAFs vs STAT1 KO + iCAFs signature scores (ULM) and iCAF-induced gene signature scores (weighted ULM) across spatial transcriptomic datasets of PDAC patients from Pei *et al*^34^. Each dot represents the median of all individual spots per patient. **b.** Violin plots showing the gene expression levels of *Stat1* adapted from Moffit *et al.* Statistical significance was calculated using a two-sided Kruskal-Wallis test. *Stat1* expression primary and normal tissues (*p* adj < 1.6 × 10^-7^), primary and metastases (*p* adj < 8.4 × 10^-9^). Normal tissue = 134, primary = 145, metastasis = 61. **c.** Median overall survival and 95% CI stratified by Stat1 expression levels (Stat1^hi^ >2.5944 n = 126 vs <= 2.5944 n = 127). **d.** Working model: In the presence of iCAFs, these release IFN-β which promotes recruitment of pSTAT1 to FOXA1-bound chromatin and drive expression of IFN-response related genes, ultimately leading to increased resistance to gemcitabine. Blocking IFN-β or knocking out STAT1 impairs iCAF-mediated transcriptional reprogramming and chemoresistance, suggesting ways to interfere with this crosstalk.

Overall, this study reveals a previously unrecognised mechanism of stromal-tumour crosstalk in PDAC, whereby PSC-derived iCAFs remodel tumour cell transcriptional and epigenetic states through paracrine signalling, particularly IFN-β. We demonstrate that iCAF-like PSCs induce pSTAT1 recruitment to enhancer regions, driving IFN-response transcriptional programme (Fig. 6d). Together, our findings position STAT1 as a key mediator of microenvironmental cues that links iCAF signalling to epigenetic reprogramming and chemotherapy resistance in PDAC.

## Discussion

In this study, we used PSC-derived engineered CAF models to recapitulate the iCAF phenotype observed in both patients and mouse models of PDAC^14–16^. Recent work demonstrated that perturbing TGF-β and IL-1 signalling *in vivo* can drive repolarisation of CAFs^23^, but this study is the first description of cell line models that stably maintain the iCAF and myCAF-like states *in vitro*.

Despite the inherent simplifications of an engineered system, our model demonstrates strong concordance with patient-derived datasets, supporting its utility as a biologically relevant system to study CAF-induced epigenetic reprogramming in pancreatic cancer cells, a hitherto under explored area of research. That said as is with any model system, there are limitations, as PSCs are not the sole precursor population contributing to CAF heterogeneity within the pancreatic tumour microenvironment^17,19,20,35^. Consequently, the engineered lines described here may be biased toward PSC-specific mechanisms, not fully capturing the complexity and diversity of CAF states observed *in vivo*.

Importantly, not all iCAF and myCAF populations arise exclusively via IL-1 or TGF-β signalling, respectively. For example, TNF signalling has also been implicated in iCAF differentiation both *in vitro* and *in vivo*^15,36^. Our model therefore does not encompass alternative or intermediate CAF states, nor the extensive plasticity that characterises these populations in patients. Future establishment of engineered models representing additional CAF phenotypes may enable broader investigation into both unique and overlapping functions of stromal populations within the PDAC microenvironment. Similarly, generation of human counterparts of these genetically “locked” CAF lines would provide a valuable platform to interrogate tumour-stroma crosstalk across mutational burden, molecular subtypes, and stages of disease progression.

Previous studies have employed bulk and single-cell transcriptomic approaches to investigate CAF tumour cell communication within the PDAC tumour microenvironment. One such study by Guinn *et al.*^37^ demonstrated that overall CAF abundance correlates with IFN and EMT signatures in epithelial tumour cells, both *in vitro* and through analysis of patient-derived single-cell RNA-seq datasets. However, when CAF populations were further stratified into iCAF and myCAF subsets, these associations were diminished, potentially reflecting the extensive phenotypic plasticity and intermediate CAF states observed in patient tumours. In contrast, our engineered system distinctly models two major CAF states in a stable manner, which may explain why activation of the IFN pathway is uniquely and reproducibly observed following iCAF co-culture. The pronounced plasticity of CAF populations remains a major challenge for therapeutic targeting, as inhibition of one CAF subtype may induce compensatory expansion or functional adaptation of other tumour-promoting stromal populations^38^. An alternative therapeutic strategy may therefore involve targeting downstream signalling dependencies within tumour cells, such as STAT1^39^, which represent context-specific vulnerabilities induced by CAF-mediated signalling. Since a subset of fibroblasts has been shown to express STAT1, its blockade could also potentially target a subset of tumour-promoting iCAFs^15,40^.

Our work identifies STAT1 as a key mediator of iCAF-induced epigenetic and transcriptional reprogramming in pancreatic cancer cells. Previous studies have associated elevated STAT1 expression with poor prognosis in PDAC patients^41,42^. Here, we describe a previously uncharacterised mechanism through which iCAFs promote tumour progression and chemoresistance via activation of the IFN-β/STAT1 axis. Chronic type I IFN signalling has previously been linked to the emergence of chemoresistance, while CAF-derived secreted factors have independently been shown to reduce chemotherapy sensitivity in pancreatic cancer^28,43^. Our study integrates these observations by demonstrating that iCAF-derived cytokines such as IFN-β can activate type I IFN signalling in tumour cells, enhancing resistance to gemcitabine treatment. It remains to be explored how these signalling mechanisms affects the immune system more broadly, particularly T cells.

We further show that pSTAT1 co-occupies a subset of genomic loci with FOXA1. Consistent with our findings, FOXA1 has previously been reported to interact with STAT1 and STAT2, while FOXA1 overexpression inversely correlates with IFN-associated transcriptional programs^32^.These observations also align with our earlier work demonstrating that FOXA1 activity is suppressed within primary tumours yet reactivated in metastatic disease^13^. Together, these data suggest that stromal-derived inflammatory signalling may dynamically reshape lineage-associated transcriptional networks in tumour cells. Whether distinct CAF populations similarly influence epigenetic programs during metastatic progression remains an important area for future investigation.

More broadly, our study highlights epigenetic regulation as an underexplored dimension of tumour-stroma crosstalk in PDAC biology. While extensive efforts have focused on cell receptor and secretory interactions between CAFs and tumour cells, comparatively little is understood about how stromal signals might reshape the epigenetic landscape of cancer cells to promote tumour progression, plasticity, and therapeutic resistance. Our findings suggest that CAF-derived inflammatory signalling can induce stable chromatin-associated changes in tumour cells, thereby reinforcing malignant phenotypes beyond transient transcriptional responses.

These observations also raise important therapeutic implications. Combinatorial treatment strategies integrating first-line chemotherapy with epigenetic modulators may provide an opportunity to target tumour cell states that are reinforced by the microenvironment cues^44,45^. Although the present study focused specifically on iCAF-induced epigenetic reprogramming, the effects of myCAF populations on tumour cell chromatin dynamics remain largely unexplored. Recent work by Roger *et al.*^46^ has further highlighted the capacity of myCAF populations to influence transcriptional plasticity and therapeutic response.

In summary, our work establishes a tractable and biologically relevant model to study stable CAF subtype-induced influence on pancreatic cancer cells. We identify the iCAF-IFN-β-STAT1 signalling axis as an important mediator of tumour cell specific epigenetic reprogramming led chemoresistance. Collectively, our study supports the growing view that effective therapeutic strategies for PDAC will likely require approaches that simultaneously target tumour-intrinsic pathways and stromal-induced adaptive states.

## Material and methods

### Cell lines

The FC1245 pancreatic cancer cell line^47^ was derived from the KPC mouse model^48^ on the C57BL/6J background and was generated in David Tuveson’s lab. All FC1245- and PSC-derived cell lines were maintained in Dulbecco’s modified Eagle’s with high glucose and pyruvate (DMEM, 41966-029, Thermo Fisher Scientific) supplemented with 5% FBS, cultured at 37 °C in 5% CO_2_ and were routinely STR-genotyped and tested for mycoplasma contamination.

### In vitro assays

For colony formation assays, FC1245-RFP^+^ cells were seeded in 24-well plates (100 cells/well). After 7 days, colonies were fixed with 4% Formaldehyde for 15 minutes and incubated with 0.05% Crystal Violet solution for at least 30 minutes at room temperature. Colonies were then washed twice with deionized water and air-dried, before being counted using the GelCount machine and software (Oxford Optronix). For conditioned media experiments, iCAFs or myCAFs were plated in 15-cm dishes (3×10^6^ cells/plate), cultured for 48 hours and subsequently starved overnight in serum-free DMEM. This media was filtered and mixed with standard 5% DMEM (1:1 dilution) and the resulting 2.5% FBS was used to culture the cancer cells. For transwell experiments, FC1245-RFP^+^ cells were plated in 6-well plates (0.3×10^6^ cells/well). A transwell with a 0.4 μm pore polycarbonate membrane (3412, Corning) was inserted onto each well and 0.3 × 10^6^ iCAFs or myCAFs were plated on top. For cytokine blocking experiments, neutralisation antibody was added on the bottom compartment.

### ELISA

Cultures of PSCs were grown for 3 days in monolayer before the culture media were collected for ELISA, performed according to manufacturer’s instructions. ELISA kits used were: IL-1α (MLA00, R&D Systems) and TGF-β1 (BMS608-4, Thermo Fisher Scientific).

### Western blotting

For Western blotting of the PSC locked lines, cells cultured in petri dishes were harvested using a cell scraper in 300 μL of RIPA lysis buffer containing protease (4693159001, Roche) and phosphatase (4906845001, Roche) inhibitors and incubated on ice for 20 minutes. Cell debris was spun down at 13,200 rpm for 10 minutes at 4 °C and the supernatant was stored at -80°C. Protein concentrations were quantified with a Bio-Rad Bradford assay, according to manufacturer’s instructions. Standard procedures were used for Western blot analysis. Briefly, 30 μg of protein was run on 4-12% gradient SDS gels for 90 minutes at 120 V, transferred for 2 hours at 90 V, blocked in 5% BSA or 10% normal mouse serum, and detected using the appropriate primary (IL1R1, AF771, R&D Systems; TGFΒR2, AF532, R&D Systems) and HRP-conjugated secondary antibodies. Pan-ACTIN (8456, Cell Signalling) was detected as loading control. For all other Western blots, cells were washed with PBS, and harvested in RIPA buffer (89901, Thermo Fisher Scientific) supplemented with protease (11836170001, Sigma Aldrich) and phosphatase (78427, Thermo Scientific) inhibitors. Lysates were then sonicated using the Bioruptor Plus (Diagenode) for a total of 3 cycles (30 seconds ON/30 seconds OFF) for live cells and 5 cycles (30 seconds ON/30 seconds OFF) for fixed cells and centrifuged at 20,000 g to collect the supernatant. Protein concentrations were measured using the Direct Detect® Spectrometer (Sigma Aldrich). Samples were denatured for 10 minutes at 70 °C in reducing agent (11569166, Thermo Fisher Scientific). 30 μg of protein were loaded onto NuPAGE 4-12% gels (NP0321, Thermo Fisher Scientific) and run for 30 minutes at 60 V and 90 minutes at 125 V before being transferred to a low-fluorescence PVDF membrane (22860, Thermo Fisher Scientific) at 30 V for 90 minutes. The membrane was blocked with Intercept Protein-Free Blocking Buffer (927-80001, LI-COR) in TBS for 1 hour at room temperature and incubated overnight at 4 °C with primary antibodies diluted in Intercept in 0.1% TBS-Tween20 (TBS-T). The membrane was washed three times with 0.1% TBS-T before incubation with secondary antibodies (Li-Cor) for 45 minutes at room temperature in Intercept (TBS-T). Subsequently, the membrane was washed twice with 0.1% TBS-T solution and once with TBS, before being developed using the Odyssey CLx Imaging System (Li-Cor).

### Generation of PSC Iocked lines

The PSC4 and PSC5 parental cells lines were previously published^14^. For the iCAF-locked lines, IL-1α (cloned in pBABE-hygro, Addgene #1765) was overexpressed in PSC4 (G1, E8) and PSC5 (F3, C9, C3) *Tgfbr2* KO PSCs from Mucciolo *et al*^24^. For the myCAF-locked lines, TGF-β1 (cloned in pBABE-hygro) was overexpressed in PSC4 (A3, A10) and PSC5 (B11, D4, E8) *Il1r1* KO PSC lines from Biffi *et al*^15^. To note, these cell lines are hygromycin resistant (pBABE-hygro), puromycin resistant (SV40 plasmid for immortalization),), blasticidin resistant (lentiCas9-Blast plasmid for Cas9 expression) and neomycin resistant (LRNG plasmid for *Il1r1* or *Tgfbr2* KO). They are also GFP^+^ (LRNG plasmid for *Il1r1* or *Tgfbr2* KO).

### STAT1 CRISPR knockout

Two CRISPR guides (Phosphorothionate-modified sgRNA, ACTGGTCGTCCAGCTGTGAG and GATGATCATGGACATCTGTA, referred to as sg792 and sg793 respectively) were designed against Exon 4 and Exon 5 of STAT1 (ENSMUST00000070968.14). FC1245-RFP^+^ cells were electroporated (Amaxa 4D Nucleofector unit, Lonza) with 4 ug TrueCut spCas9 protein V2 (Invitrogen A36498) and 80 pmol guide RNA (Synthego), using program CM137 and P3 nucleofector solution. Single cells were seeded into 96 well plates by FACS 3 days post electroporation into 50% conditioned media to derive individual clones. A cell pellet from each pool was taken 3 days and 10 days post electroporation, as well as from a number of clones, and genomic (g)DNA was extracted (Qiagen, DNeasy blood and tissue kit, 69506). Exon 4 and Exon 5 of STAT1 were amplified by PCR (Q5 High-Fidelity DNA polymerase, NEB, cat#M0491S) using Exon 4 forward primer (3841): ACGTGACTATCACCAGTGGC and reverse primer (3842): CAGTGAGCGTGCACTCAGTA for screening for sg792 editing, and Exon 5 forward primer (3843): ATGACCCACAGTTATTCTGCGAT and reverse primer (3844): TCTTACACCCACGTGCCATTT for screening for sg793 editing. Amplicons were subjected to Sanger sequencing and analysed using Synthego ICE web tool to calculate the percent editing in a pool and to determine the genotype of specific clones. Two STAT1 KO clones and two Cas9 controls were pooled together.

### Animals

#### Ethical

All animal studies were conducted in accordance with UK Home Office regulations under project license P1919999D or PP6681429 and approved by the Cancer Research UK (CRUK) Cambridge Institute Animal Welfare and Ethical Review Board. Female and male NSG mice (Charles River Laboratories) were housed in individually ventilated cages, with wood chip bedding and nestled with environmental enrichment (cardboard fun tunnels and chew blocks) under a 12-hour light/dark cycle at 21 ± 2 °C and 55 ± 10% humidity.

#### Orthotopic implantation

Pancreatic orthotopic injections were conducted as previously described^49^. Cells were single cell suspended in 50% Matrigel in PBS and injected into the tail of each pancreas. For the co-injection study, 50 μL containing 5,000 FC1245-RFP^+^ cells were injected alone or co-injected with iCAFs and/or myCAFs at a ratio of 1:4. For the STAT1 KO study, 30 μL containing 5,000 Cas9-Control or STAT1 KO cells were co-injected with iCAFs at a ratio of 1:10. From day 7 post-implantation onwards, mice were intraperitoneally injected with 120 mg/kg gemcitabine or vehicle (Vetivex saline solution) twice a week. Mice were allowed to recover from surgery for a period of one week prior to weekly abdominal palpation to determine the presence of any pancreatic masses. Once confirmed, mice were closely monitored for clinical signs and morbidity, evidenced by decreased grooming, hunched posture and reduced movement. Disease progression was monitored until clinical endpoint.

#### ATAC-sequencing

ATAC sequencing for fixed cells (InTAC-seq) was adapted from a previously described protocol^26^. Briefly, 1×10^6^ FC1245-RFP^+^ cells were cultured alone or together with 3×10^6^ iCAFs or myCAFs for 48 h. Cells were then fixed with 1.6% formaldehyde in PBS for 5 min, then quenched with an equal volume of 1×eBioscience permeabilization buffer (Thermofisher Scientific, 00-8333-56). Cells were centrifuged for 5 minutes at 600 g and incubated with the permeabilization buffer for 30 mins at room temperature. 50,000 FC1245-RFP^+^ cells were sorted using the FACSMelody Cell sorter (BD Biosciences) and used for downstream processing. Cells were spun down at 600 g for 5 mins and resuspended in transposition mix containing 1×TD buffer, 0.1% NP40, 0.01% digitonin, and Tn5. Cells were incubated for 1 hour at 37 °C with 1200 rpm shaking. 2×reverse crosslinking buffer (2% SDS, 0.2 mg/mL proteinase K, and 100 mM N,N-Dimethylethylenediamine, pH 6.5 [Sigma Aldrich D158003]) was added at equal volume to transposed cells and reversal of crosslinks was performed at 37 °C overnight with 600 rpm shaking. DNA was purified using the Zymo DNA Clean and Concentrator-5 Kit (D4014), and size selection was performed using Ampure XP beads. ATAC-seq libraries were generated using the Nextera XT Index kit (24 indexes, FC-131-1001) or the Illumina DNA-RNA UD indexes and sequenced with 50 bp paired end reads using NovaSeq 2000 or NovaSeq X (Illumina).

#### ChIP-sequencing

1×10^6^ FC1245-RFP^+^ cells were plated alone or together with 3×10^6^ iCAFs or 3×10^6^ myCAFs. Cells were double crosslinked 48 hours later, firstly with 2 mM disuccinimidyl glutarate (DSG) for 20 minutes and then with 1% formaldehyde for 10 minutes before crosslinking was quenched with 0.1 M glycine for 5 min. Cells were then washed twice in cold PBS and harvested in cold PBS containing protease and phosphatase inhibitors. For H3K27ac and H3K4me1 ChIP-seq, crosslinking was performed in suspension and FC1245-RFP+ cells were sorted and stored at -80 °C before proceeding with cell lysis. Crosslinked samples were lysed in lysis buffer 1 (LB1, 50 mM Hepes–KOH, pH 7.5, 140 mM NaCl, 1 mM EDTA, 10% Glycerol, 0.5% NP-40/Igepal CA-630, 0.25% Triton X-100), centrifuged at 4,000 rpm for 5 minutes at 4 °C and lysed in lysis buffer 2 (LB2, 10 mM Tris–HCL, pH8.0, 200 mM NaCl, 1 mM EDTA, 0.5 mM EGTA) before resuspension in lysis buffer 3 (LB3, 10 mM Tris–HCl, pH 8, 100 mM NaCl, 1 mM EDTA, 0.5 mM EGTA, 0.1% Na–Deoxycholate, 0.5% N-lauroylsarcosine). Chromatin was sonicated using the Bioruptor Plus (Diagenode) for 20 cycles (30 seconds ON/30 seconds OFF) to generate DNA fragments of around 100-800 bp. Beads were pre-bound with antibody overnight at 4 °C, for which 2.5 μg of antibody and 25 μL of Protein A or G Dynabeads (Invitrogen) were used per each 15 cm plate. Two 15-cm plates were used per IP. Chromatin and beads were incubated overnight at 4 °C rotating at 12 rpm. The following day, beads were washed 6 times in cold RIPA buffer (50mM HEPES pH 7.6, 1mM EDTA, 0.7% Na-deoxycholate, 1% NP-40, 0.5M LiCl) and once in cold Tris-EDTA (10mM Tris-HCl, 1mM EDTA). Samples were incubated in elution buffer (50mM TrisHCl, pH8, 10mM EDTA, 1% SDS) for 18-24 hours at 65 °C for reversal of crosslinks. Eluted DNA was treated with 20 ng/ml RNase A (Invitrogen, AM2271) for 1 hour at 37 °C and with 200 ng/ml proteinase K (ThermoFisher Scientific, 25530049) for 2 hours at 55 °C before DNA was extracted using phenol-chloroform. Size selection was performed with AMPure XP Beads (Beckman CoulterTM). Library preparation was performed using the SMARTer ThruPlex DNA-seq kit (Takara R400676), and samples were sequenced using the NovaSeq 2000, 6000 or NovaSeq X 50-bp paired-end sequencing (Illumina).

#### ATAC-sequencing and ChIP-sequencing analyses

Reads were mapped to the mm10 genome using bowtie2 version 2.2.6. Aligned reads with mapping quality <5 were filtered out. Read alignments from all replicates were combined into a single library and peaks were called with MACS2 version 2.1.1.2016 using input sequences as background control. The peaks that absorbed statistically significant tag density from all replicates (q ≤ 1 × 10^-3^) were selected for downstream analysis. Differential binding analysis (DiffBind) was performed as described previously^27^. Motifs were identified among tag-enriched sequences by using Meme v.4.9.1. For visualization of tag density and signal distribution heatmap, the read coverage in a window of a region ± 2.5 or 5 kb flanking the tag midpoint was generated using a bin size 1/100 of window length.

#### RNA-sequencing

For the PSC-derived cell lines, cells were pelleted and lysed in TRIzol Reagent (15596-018, Thermo Fisher Scientific) and RNA was extracted with an Invitrogen PureLink RNA Mini Kit (12183018A, Thermo Fisher Scientific), according to manufacturer’s instructions. For all other cell lines, cells were fixed with 1% paraformaldehyde and either sorted from co-culture or directly harvested and stored at -80 °C. RNA was extracted from using the Maxwell^®^ RSC simplyRNA Tissue Kit (AS1340, Promega). RNA concentration was assessed using Qubit and RNA quality was evaluated using the RNA Integrity Number (RINe) from Tapestation 4200 system, and the TruSeq stranded mRNA library preparation kit (20020594, Illumina) was used to prepare sequencing libraries. RNA sequencing was performed using NovaSeq 2000, NovaSeq 6000 or NovaSeqX 50-bp paired-end sequencing (Illumina).

#### RNA-sequencing analysis

All sequencing was carried out by the Genomics Core Facility (CRUK-CI).

For RNA-sequencing analysis of PSC-derived cell lines, FASTQ files were aligned, and the expression levels of each transcript were quantified using Salmon (v1.4.0)^50^ with the annotation from ENSEMBL (GRCh38 release 98) with recommended settings. Transcript-level expression was loaded and summarized to the gene level by using tximport^51^. Differential gene expression was performed using DESeq2 v2^52^ by applying lfcshrink function^53^. The principal components for variance-stabilized data were estimated using plotPCA function, available in DESeq, and ggplot2^54^. Genes with adjusted *p* value < 0.05 were selected as significantly differentiated between conditions. Following differential gene expression, genes were pre-ranked based on the negative logarithmic p-value and the sign of the log_2_ fold change. GSEA was performed using clusterprofiler^55^ against the Hallmark, Reactome, and C2 canonical pathway collection (C2.cp.v5.1) downloaded from the Molecular Signatures Database (MSigDB)^56^. GSEA plots were plotted using enrichplot R package^57^. For RNA-sequencing analysis of the FC1245-RFP^+^ parental, STAT1 KO and Cas9 control cell lines, reads were mapped to the mm10 reference genome using bowtie2 version 2.2.6. Read counting was performed using featureCounts program of Subread software package version 2.0.4. Differential gene-expression analysis used the DESeq2 v2 workflow.

#### qPLEX-RIME

Chromatin preparation and immunoprecipitation were performed as described for ChIP-seq. After overnight incubation with the chromatin, beads were washed 10 times in cold RIPA buffer (50 mM HEPES pH 7.6, 1 mM EDTA, 0.7% Na-deoxycholate, 1% NP-40, 0.5 M LiCl) and twice in 100 mM ammonium hydrogen carbonate (AMBIC). Washed beads were frozen at -80 °C until further processing. RIME sample preparation and analysis was performed by the Proteomics Core Facility (CRUK-CI) as previously described^30,31^. Tryptic digestion of bead-bound protein was performed by addition of 15 ng/μL trypsin (Pierce) in 100 mM AMBIC followed by overnight incubation at 37 °C. A second digestion step was performed the next day for 4 h. After tryptic digestion, supernatant was acidified by adding 2 μL 5% formic acid. Digested samples were cleaned using Ultra-Micro C18 Spin Columns (Harvard Apparatus) according to manufacturer’s instructions. For quantitative RIME (qPLEX-RIME), digested, cleaned peptide samples were dried using a speedvac, reconstituted in 25 μL 0.1 M TEAB and labelled using TMTpro-18plex reagents (Thermo Fisher) following a randomised design. The peptide mixture was fractionated with Reversed-Phase cartridges at high pH (Pierce). Nine fractions were collected using different elution solutions in the range of 5-50% ACN and dried. Each fraction was reconstituted in 0.1% formic acid for and analysed on a Dionex UltiMate 3000 UHPLC system coupled with the nano-ESI Fusion-Lumos (Thermo Scientific). The Proteome Discoverer 2.4 (ThermoFisher Scientific) was used for the processing of raw MS files. The SequestHT search engine was used and all the spectra searched against the Uniprot Mus musculus FASTA database (taxon ID 10090 - Version Jan 2022). All searches were performed using a static modification of TMTpro (+304.207 Da) at any N-terminus and lysines and Carbamidomethyl at Cysteines (+57.021 Da). Methionine oxidation (+15.9949 Da), and Deamidation on Asparagine and Glutamine (+0.984) were included as dynamic modifications. Mass spectra were searched using precursor ion tolerance 20 ppm and fragment ion tolerance 0.5 Da. For peptide confidence, 1% FDR was applied, and peptides uniquely matched to a protein were used for quantification. For downstream data analysis, the R package qPLEXanalyzer was used^31^. Statistical analyses were performed using a limma-based differential test.

## Patient data analysis

### Spatial transcriptomics

Spatial transcriptomic data from Pei *et al.*^34^ were obtained from the Sequence Read Archive (SRA accession: SRP525961). The loupe browser (10x Genomics) was used to annotate tumour and stroma regions. Raw sequencing data from 13 patient samples were processed using Space Ranger v4.0.1 (10x Genomics) with default parameters to generate spatial gene expression matrices and align spots to tissue histology images. Processed spatial transcriptomic data were analysed using the Seurat package and normalised using the sctransform package. Gene signature scores for iCAF presence and iCAF-induced signatures were calculated using the decoupleR package with univariate linear model (ulm). The iCAF signature was previously described by Biffi *et al.*^15^ and Dimi *et al.* The iCAF-induced signature consisted of differentially expressed genes upregulated in FC1245-RFP^+^ cells after 48-hour co-culture with iCAFs (log_2_FC > 1, *p* < 0.05). Correlation between iCAF presence scores and iCAF-induced signature scores was assessed across all spatial spots using Pearson correlation. Data visualization was performed using ggplot2, tidyverse, and patchwork packages. Spatial data were visualized with Loupe Browser (10x Genomics).

### COMPASS trial

A total of 253 patients with RNA-sequencing and survival data from the COMPASS trial (NCT02750657)^58^ were analysed. The COMPASS trial was a prospective study designed to explore biomarkers of chemotherapy response and was approved by the University Health Network Research Ethics Board (REB:15-9596). Patients were stratified based on *STAT1* expression levels. Overall survival (OS) was calculated from the date of enrolment. Kaplan-Meier survival curves were generated and compared using the log-rank (Mantel-Cox) test. Hazard ratios and 95% confidence intervals were calculated using Cox proportional hazards regression models.

### Statistical analysis

GraphPad Prism and custom R or Python scripts were used for graphical representation of data. For experiments comparing iCAF- and myCAF-locked PSCs, statistical analyses were performed using non-parametric Mann-Whitney test. For three group comparisons *in vitro*, parametric one-way ANOVA or an extra sum-of-squares F test to test differences between growth curves were used. For the RIME (proteomics) dataset, a limma-based analysis was preferred to shrink individual protein variances toward a global average. For *in vivo* and patient survival analysis, we displayed the survival probability of each group as a function of time by means of Kaplan-Meier plots and used the log-rank (Mantel Cox) test to test the null hypothesis that there is no difference between the populations in the probability of death at any time point. All statistical details of experiments are specified in the respective figure legends or method sections, including number of replicates and how significance was defined.

## Material availability

Cell lines generated in this study are available upon request. CAF-locked lines are available upon request from G.B. lab.

## Acknowledgements

We thank the core facilities at Cancer Research UK (Biological Research Unit, Genomics, Proteomics, Histopathology, Bioinformatics, Research Instrumentation and Flow Cytometry) for their consistent support and advice. We would like to acknowledge the funding support from:

1. University of Cambridge, Cancer Research UK Cambridge Institute core funds (Grant number: G101107) – J.S.C.
2. Pancreatic Cancer UK Future Leaders Academy (Grant number: FLF_2021_05) – G.B., J.S.C.
3. Pancreatic Cancer Research Fund (Grant number: G110303) – S.V.R., J.S.C.
4. University of Cambridge Cancer Research UK Cambridge Institute core funds (Grant number: A27463) – G.B.
5. CRUK-NCI Cancer Grand Challenge (CANCAN team) – G.B.
6. UKRI Future Leaders Fellowship – G.B.

The Fusion Lumos Orbitrap mass spectrometer was purchased with the support from a Wellcome Trust Multi-user Equipment Grant (Grant # 108467/Z/15/Z).

## Author Contributions

Conceptualisation, S.V.R., C.P., G.B., and J.S.C.;

Investigation and data collection, C.P., M.W., P.S.W.C., L.Y., E.M., Y.C., S.K., M.C., S.P.T., V.N.R.F., E.K.P., C.D., A.R.;

Formal analysis, C.P., I.C., M.J., A.R.E., K.K., C.S.R.C.;

Visualisation, C.P., I.C., M.J.; A.R.E.,

Funding acquisition and supervision, S.V.R., G.B. and J.S.C.;

Writing – original draft, C.P., S.V.R., G.B., J.S.C.;

Writing – review & editing, all authors.

## Extended Figure legends

**Extended Figure 1.**
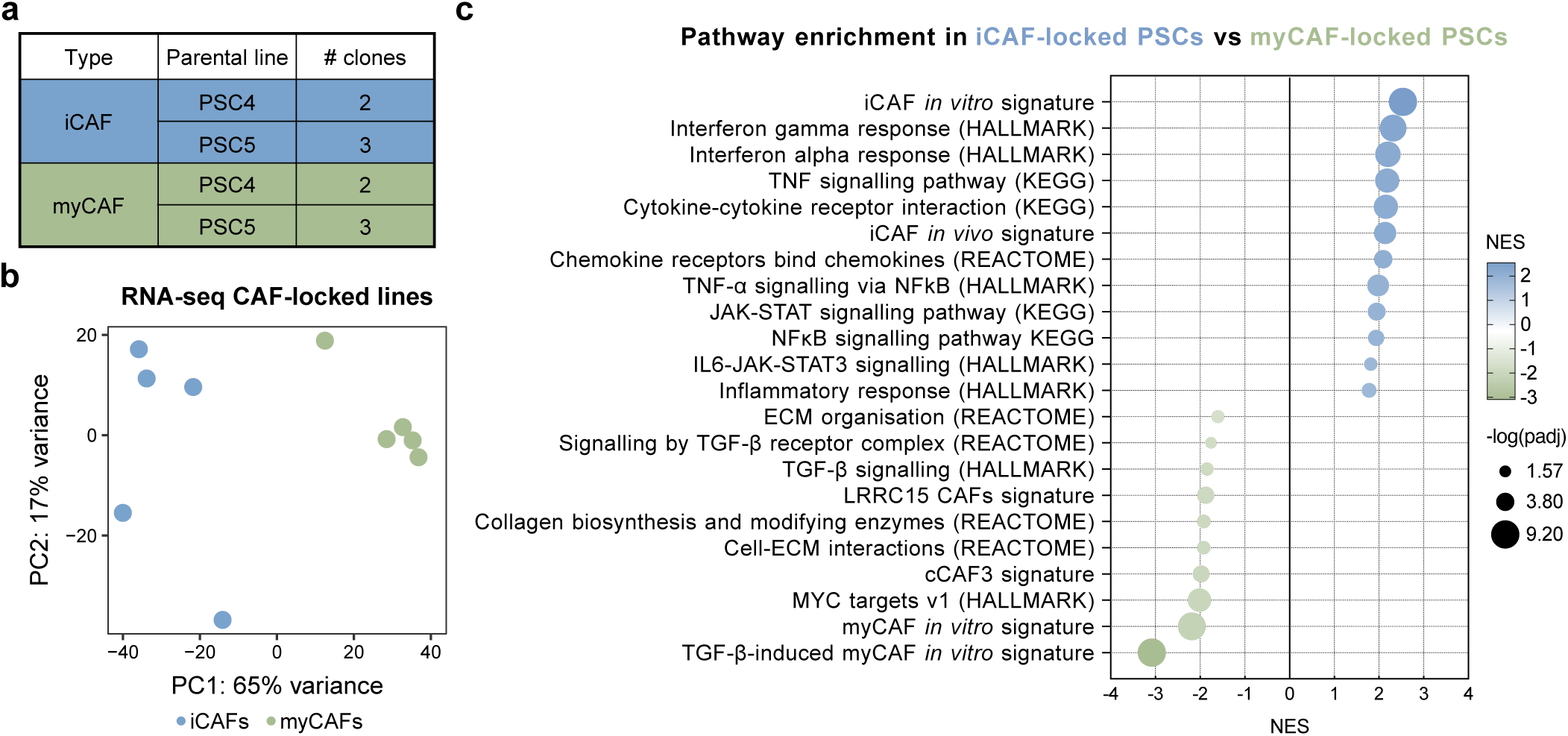
**a.** Table showing the clones of CAF-locked PSCs analysed by RNA-sequencing. **b.** Principal component analysis plot of CAF-locked PSC transcriptomes. **c.** GSE analysis comparing iCAF-locked to myCAF-locked PSCs. NES, normalised enrichment score. padj, adjusted *p* value. ECM, extracellular matrix. The TGF-β-induced myCAF *in vitro*, LRRC15^+^ and cCAF3^+^ signatures are from Mucciolo *et al*^24^. The iCAF and myCAF *in vitro* signatures are from Öhlund *et al*^14^. The iCAF *in vivo* signature is from Elyada *et al*^16^.

**Extended Figure 2.**
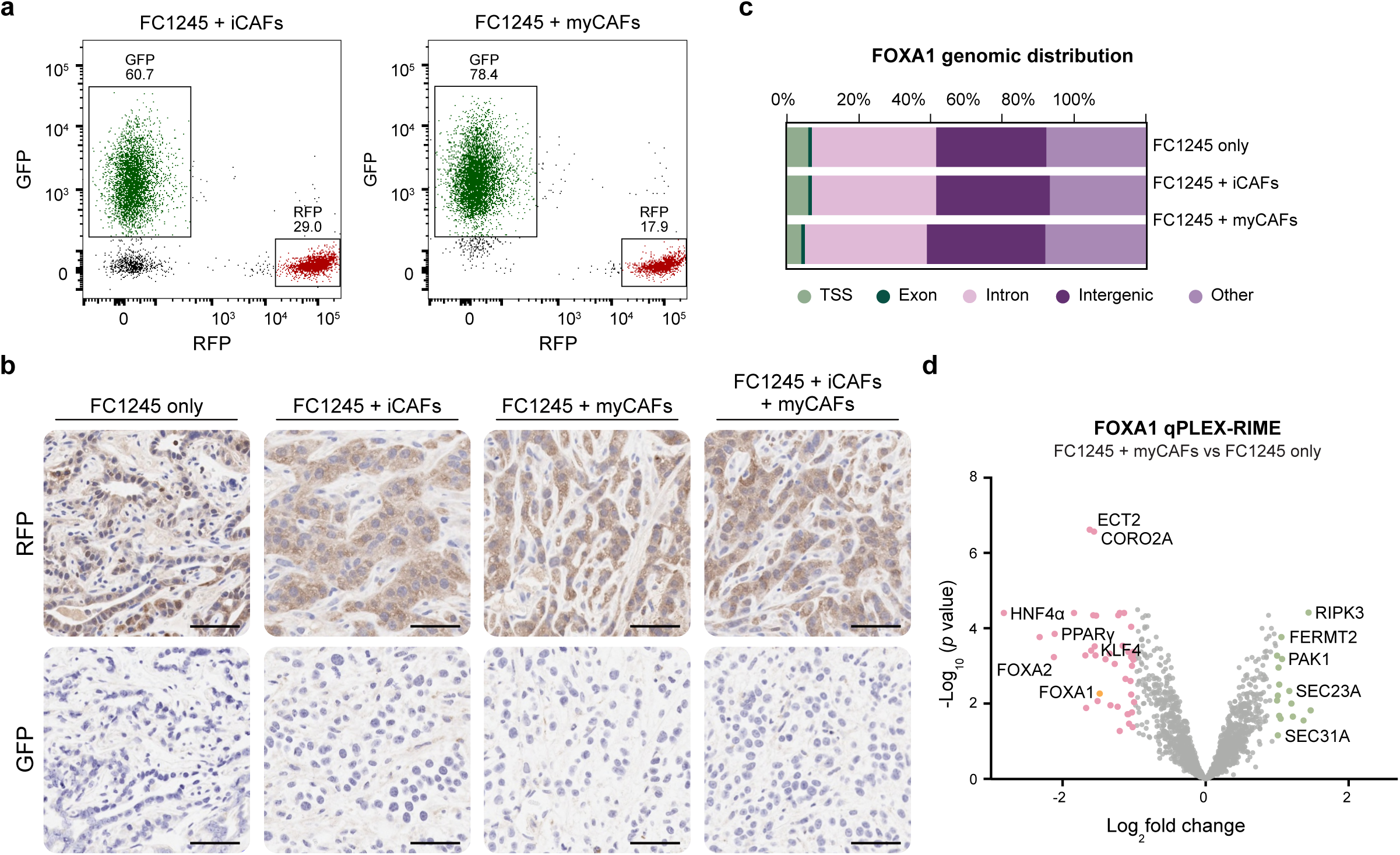
**a.** Representative FACS plots showing the sorting strategy used sort FC1245-RFP^+^ cancer cells and iCAFs-GFP*^+^* (left panel) or myCAFs-GFP+ (right panel) prior to orthotopic injection. **b.** Representative images of GFP and RFP immunohistochemistry (IHC) staining of FC1245 only, FC1245 + iCAFs, FC1245 + myCAFs and FC1245 + iCAFs + myCAFs orthotopic tumours. Scale bar is 50 µm. **c.** Genomic feature distribution of FOXA1 in FC1245-RFP^+^ cells cultured alone (FC1245 only), or co-culture with iCAFs (FC1245 + iCAFs) or myCAFs (FC1245 + myCAFs), n=3. **d.** Volcano plot of FOXA1 qPLEX-RIME data representing FOXA1 interactors differentially regulated after co-culture with myCAFs, n=6. Proteins with log_2_fold change > 1 and *p* value < 0.05 are highlighted in blue (significantly upregulated by iCAFs) and proteins with log_2_fold change < -1 and *p* value < 0.05 are highlighted in pink (significantly upregulated in monoculture). Statistical significance was calculated using a limma-based differential analysis.

**Extended Figure 3.**
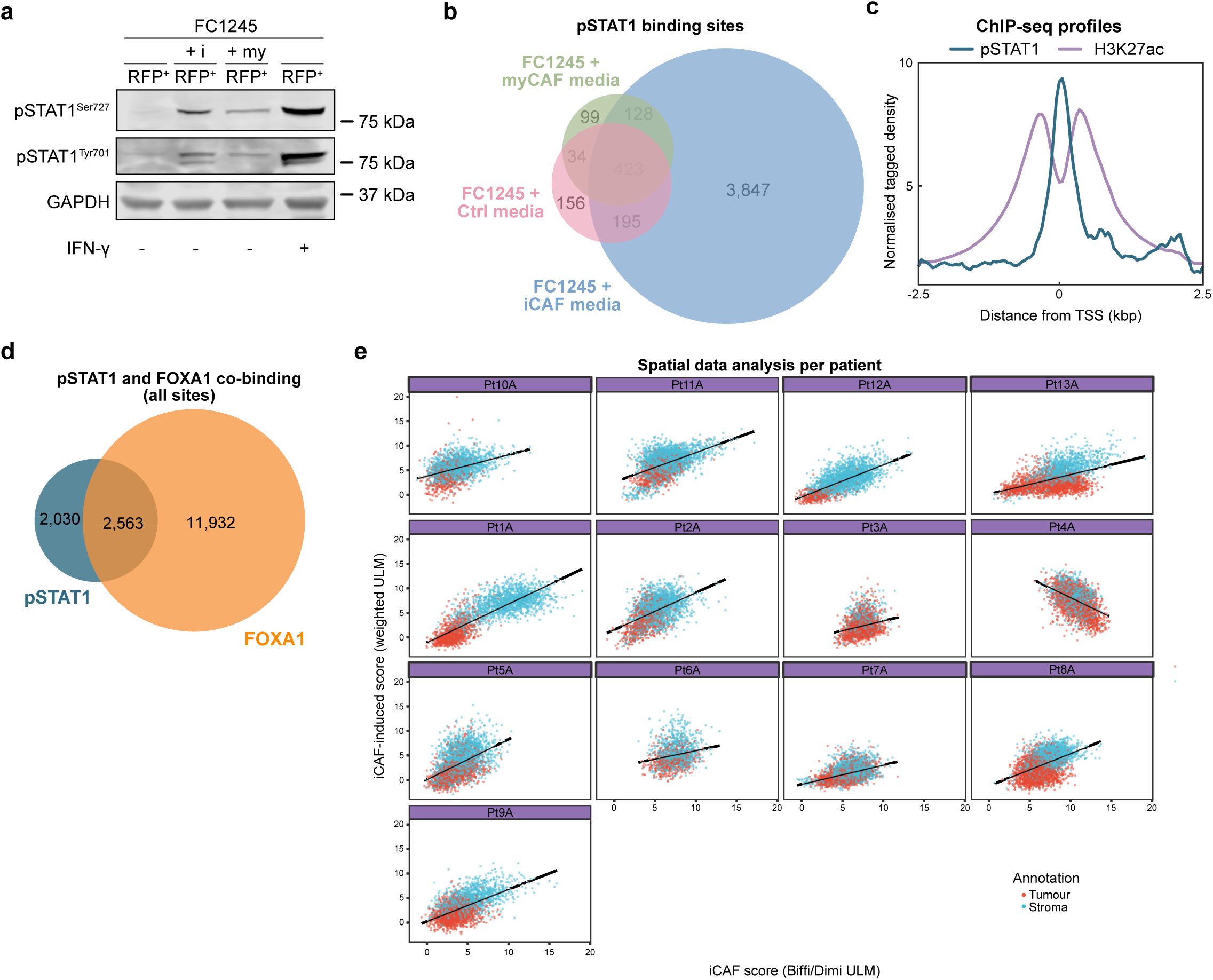
**a**. Western blotting of pSTAT1^Ser727^ and pSTAT1^Tyr701^ in FC1245 (RFP^+^) after monoculture, co-culture with iCAFs or myCAFs or stimulation with 20 ng/mL of IFN-γ for 1 hour as a positive control (right panel). GAPDH was used as a loading control. **b.** Venn diagram indicating the overlap of pSTAT1 binding sites in FC1245-RFP^+^ cells cultured in iCAF- or myCAF-conditioned media, or in control media, n=4. **c.** ChIP-seq profiles of pSTAT1 and H3K27ac distribution in FC1245-RFP^+^ cells cultured in iCAF-conditioned media. **d.** Venn diagram indicating the overlap between total pSTAT1^Tyr701^ and FOXA1 binding sites in FC1245-RFP^+^ cells cultured in iCAF-conditioned media or together with iCAFs, respectively. **e.** Scatter plots showing the correlation between iCAF signature scores (indicating iCAF presence) and iCAF-induced gene signature scores in all spots from each individual patient using spatial transcriptomic datasets from Pei *et al*^34^.

**Extended Figure 4.**
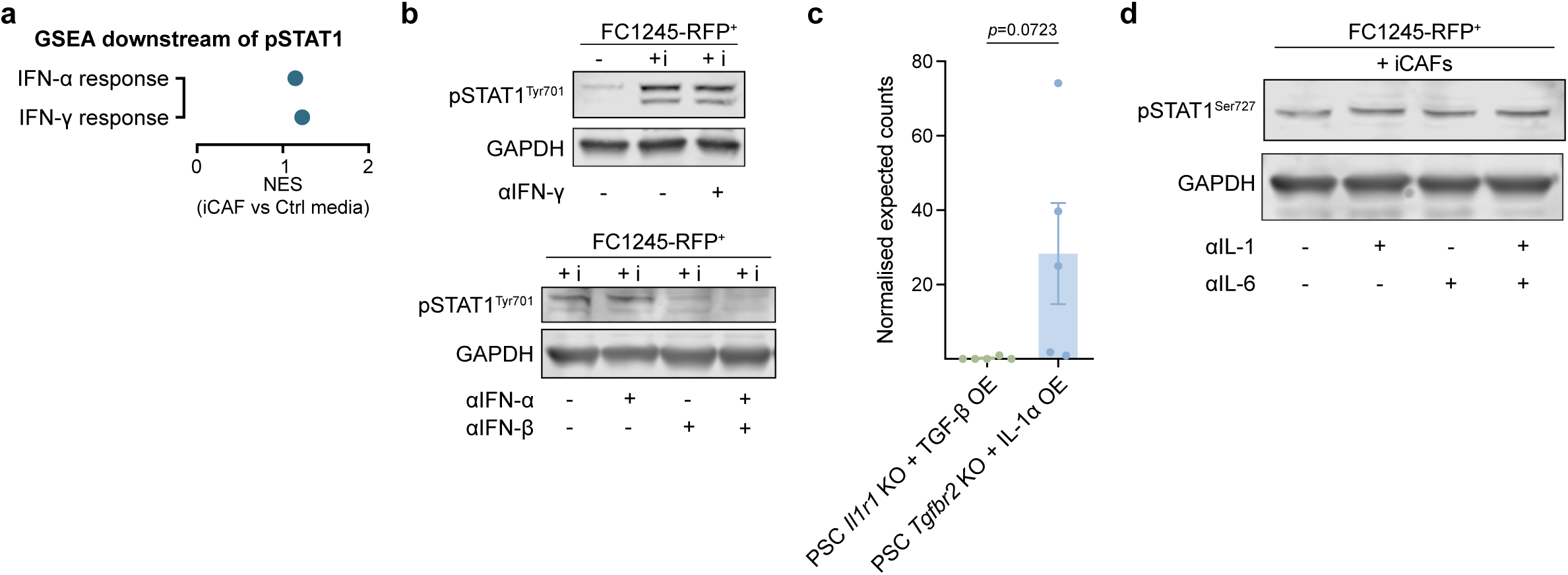
**a.** GSE analysis of genes located downstream of promoters that pSTAT1 binds to in FC1245-RFP^+^ + iCAF media conditions. **b.** Western blotting of pSTAT1T^yr701^ in FC1245-RFP^+^ cells cultured on their own (-) or separated from iCAFs by a transwell (+i) and treated with 10 µg/mL of anti-IFN-γ neutralising antibody for 48 hours (top panel) or with 25 µg/mL of anti-IFN-α or/and 2,000 neutralising units (NU)/mL of anti-IFN-β for 48 hours (bottom panel). **c.** Normalised expected counts of *Ifnb* in myCAF-locked (green) or iCAF-locked (blue) PSCs. Statistical significance was calculated using a two-tailed unpaired t-test. **d.** Western blotting of pSTAT1^Ser727^ in FC1245-RFP^+^ cells separated from iCAFs by a transwell and treated with 5 µg/mL of anti-IL-1α or/and 10 µg/mL of anti-IL-6 neutralising antibodies for 48 hours.

**Extended Figure 5.**
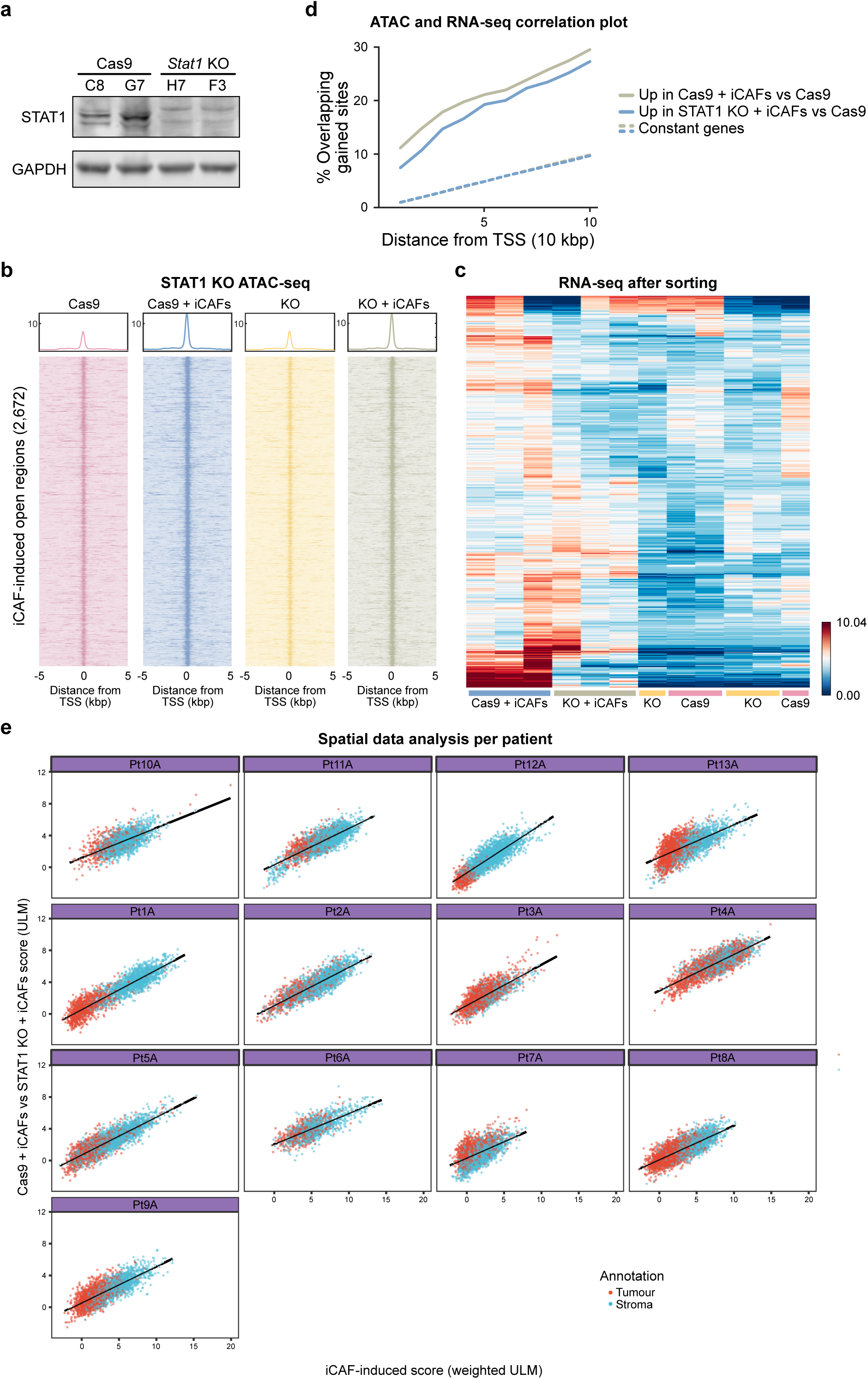
**a.** Western blotting of STAT1 and GAPDH in the Cas9-only and STAT1 KO clones that were merged for ATAC-seq and *in vivo* experiments. **b.** Heatmaps of ATAC-seq signal at iCAF-induced *DiffBind* open regions (2,672 sites) gained in STAT1 KO or Cas9 controls before or after co-culture with iCAFs, n=3. **c.** Differential gene expression analysis of STAT1 KO or Cas9 cells cultured alone or sorted from co-culture with iCAFs, n=3. **d.** Enrichment of the iCAF-induced open regions (2,672 sites) in the vicinity of genes upregulated in STAT1 KO or Cas9 only clones after co-culture with iCAFs (compared to Cas9 alone). **e.** Scatter plots showing the correlation between Cas9+iCAFs vs STAT1 KO+iCAFs signatures score and iCAF-induced signature scores in all spots from each individual patient using spatial transcriptomic datasets from Pei *et al*^34^.

## Notes

### Competing Interest Statement

The authors have declared no competing interest.

